# Loss of MITF activity leads to emergent cell states from the melanocyte stem cell lineage

**DOI:** 10.64898/2025.12.23.695681

**Authors:** Alessandro Brombin, Stephanie MacMaster, Jana Travnickova, Cameron Wyatt, Hannah Brunsdon, Emma Ramsey, Hong Nhung Vu, Eirikur Steingrimsson, Colin Kenny, Tamir Chandra, E. Elizabeth Patton

## Abstract

How embryonic cells generate large clones of cells in the adult represents a fundamental question in biology. Here using melanocyte stem cells (McSCs) in the zebrafish as a model we explore the function of the master melanocyte transcription factor (MITF) in safeguarding McSCs in embryonic development and their potential to pigment large clones in the adult. MITF is well known is for its role in the specification of melanoblasts from the neural crest (NC) and their differentiation into melanocytes, yet little is known about how this activity shapes the stem cell lineages. Here, we use live imaging coupled with single-cell transcriptomics and lineage tracing to show that MITF (*mitfa* in zebrafish) protects the melanocyte stem cell (McSC) fate in zebrafish. Utilizing a temperature sensitive *mitfa^vc7^* mutant, we show that loss of Mitfa activity leads to a surprising premature and aberrant expansion of McSC progeny at the niche during embryogenesis, coupled with novel emergent transcriptional cell states. Linage tracing of McSCs from the embryonic to juvenile stages reveals Mitfa activity is subsequently required in regeneration by Schwann cell-like and melanocyte stem cell progenitors that serve as a reservoir for fast-responding pigment progenitors. Thus, the impact of Mitfa loss on the melanocyte lineage is cell-state and stage-specific. The emergent cell states resulting from *mitfa* loss may have important implications for understanding how reduced MITF activity contributes to human genetic disease and melanoma.

## Introduction

Melanocyte stem cells (McSCs) give rise to large clones of pigment cells on the adult zebrafish skin, yet little is understood about how these embryonic cell types give rise to the complex pigmentation of the adult zebrafish. Melanocytes evolved to produce melanin, a UV-absorbing polymer that is responsible for the colour spectrum of our skin, hair, and eyes, and contributes to the array of colours and patterns in the animal kingdom (Kaelin and Barsh 2013; Patterson and Parichy 2019; Kratochwil and Mallarino 2023; Brombin and Patton 2024). In less well understood physiology, melanocytes can also be found throughout the body in immune, auditory, heart and neuroendocrine systems (Yajima and Larue 2008; Levin et al. 2009; Plonka et al. 2009; Kaucka et al. 2021). The embryological origins of these diverse melanocytes and the territories that they subsequently populate in the adult represent a wide knowledge gap that is directly relevant to melanocyte patterns and melanocytic disease.

In all animals, melanocyte differentiation is dependent upon the highly conserved melanocyte inducing transcription factor (MITF) (Goding and Arnheiter 2019). In both melanoma and melanocyte development, MITF activity is a tightly controlled molecular “rheostat” leading to divergent cellular outcomes. In general terms, high MITF activity promotes pigmentation and cell cycle arrest, moderate MITF activity promotes proliferation, and low-to-no MITF activity leads to migration, stem like states or cell death, depending on the cellular context (McGill et al. 2002; Carreira et al. 2006; Taylor et al. 2011; Muller et al. 2014; Kawakami and Fisher 2017; Rambow et al. 2018; Goding and Arnheiter 2019; Nguyen and Fisher 2019; Rambow et al. 2019; Travnickova et al. 2019; Travnickova et al. 2022; Centeno et al. 2023). Further layers of regulation to MITF activity are added by post-translational modifications, including acetylation and phosphorylation that fine-tune the intensity, specificity and selectivity of MITF binding patterns in the genome (Leclerc et al. 2017; Ngeow et al. 2018; Louphrasitthiphol et al. 2020; Louphrasitthiphol et al. 2023; Vu et al. 2024). MITF post-translational modifications can have direct relevance for disease, as illustrated by germline mutations in the sumoylation motif of MITF that are associated with sporadic and familial melanoma (Bertolotto et al. 2011; Yokoyama et al. 2011; Bonet et al. 2017). In addition to a well-established MITF transcriptional activator role, MITF has a less well understood repressor function in melanoma cells, that includes repression of extracellular matrix gene expression and oxidative phosphorylation genes (Strub et al. 2011; Laurette et al. 2015; Mort et al. 2015; Berico et al. 2021; Dilshat et al. 2021; Dias et al. 2025). Thus, MITF is a powerful transcriptional activator, but also has potent gene repressor functions in melanoma. Additionally, in MITF-high conditions, MITF can inhibit gene expression indirectly by limiting the stability and nuclear activity of TFE3 in melanoma (Chang et al. 2025).

In zebrafish development, melanocytes originate directly from the NC, along with two other pigment cell types, the silver iridophores and the yellow xanthophores, to patten the zebrafish embryo (Kelsh et al. 1996; Lister et al. 1999; Singh and Nusslein-Volhard 2015; Irion et al. 2016; Patterson and Parichy 2019). During NC migration, in addition to the direct differentiation of embryonic melanocytes, a subpopulation of *mitfa+* cells migrate to the prospective dorsal root ganglia (DRG) to become melanocyte stem cells (McSCs) **(Figure 1A).** (Budi et al. 2008; Hultman et al. 2009; Budi et al. 2011; Dooley et al. 2013; Singh et al. 2016; Brombin et al. 2022). These ERB-signalling dependent cells contribute to a few of the melanocytes in the embryonic lateral stripe, but are otherwise mostly quiescent until they are called upon to contribute to <u>m</u>elanocytes, iridophores, <u>x</u>anthophores, and/or nerves, in the adult skin during growth or regeneration (and thereby named MIX+ cells (Hultman et al. 2009; Dooley et al. 2013; Singh et al. 2016; Johansson et al. 2020; Brombin et al. 2022; Brunsdon et al. 2022). We have previously shown that McSC transcription and ALDH2 metabolic activity are tightly controlled in melanocyte regeneration (Johansson et al. 2020; Brunsdon et al. 2022). Further, we showed that *tfap2b* is a functional marker of adult McSCs in embryos, and that *tfap2b+* cells contribute to large, multi-cellular clones in the adult skin (Brombin et al. 2022).

**Figure 1:**
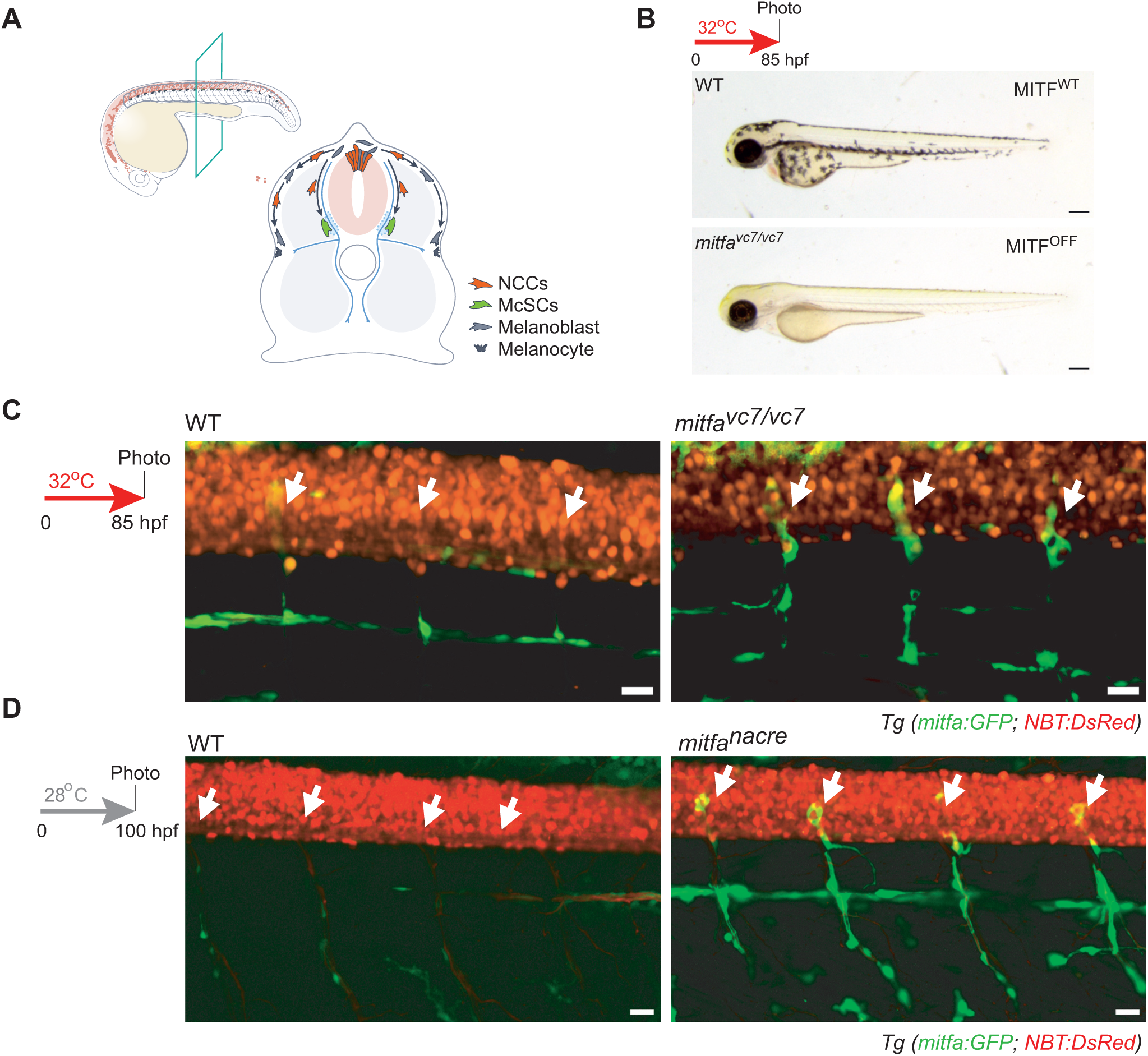
McSC progeny expand in absence of MITF activity. **A.** McSCs in the zebrafish embryo. Schematic of a zebrafish embryo (approximately 24 hours post fertilization; hpf) and transverse section showing melanoblasts and melanocytes originating from the neural crest cells (NCCs). Melanocyte stem cells (McSCs) are established at the site of the prospective dorsal root ganglia (DRG); dashed blue lines. These cells become quiescent once at the niche and produce melanocytes at later stages of growth or during regeneration. Illustrations by Uta Mackensen. **B.** Mitfa activity can be modulated in the *mitfa^vc7/vc7^* temperature sensitive mutant. Images of WT (top) and *mitfa^vc7/vc7^* (bottom; no visible embryonic melanocytes) held at non-permissive temperature of 32°C for 72h to reach the 85 hpf developmental equivalent. Scale bar: 200 μm. **C.** McSCs expand in *mitfa^vc7^* mutant lines. Confocal stacks (30 µm) of *Tg(mitfa:GFP; NBT:DsRed)* embryos either in WT (left) or *mitfa^vc7/vc7^*(right) mutants. Fish were raised at 32°C for 72h (85 hpf equivalent). McSCs (white arrows) turn off *mitfa* expression (no *mitfa:GFP* expression) in WT embryos, with a few embryonic GFP+ melanoblasts migrating along nerves to pattern the embryo. McSCs and progeny are expanded in the absence of Mitfa activity in *mitfa^vc7/vc7^* mutants. Standard deviation projection. Scale bar: 20 μm. **D.** McSCs expand in *mitfa^nacre^* mutant lines. Confocal stacks (30 µm) of *Tg(mitfa:GFP; NBT:DsRed)* embryos either on WT (left) or *nacre* (constitutive *mitfa* mutant; right) background. Fish were raised at 28°C until 100 hpf. McSCs (white arrows) turn off *mitfa* expression (no *mitfa:GFP* expression) in WT embryos while McSC progeny are expanded in the *nacre* mutant. Standard deviation projection. Scale bar: 20 μm.

Over fifteen years ago, Johnson, Lister and colleagues showed that despite the requirement of Mitfa for melanocyte differentiation, Mitfa activity is not required for the establishment of the McSC at the niche (Johnson et al. 2011; Zeng et al. 2015). Using a temperature sensitive splice site mutation in *mitfa*, called *mitfa^vc7/vc7^*, that is controlled by changing the water temperature, they showed that Mitfa is essential for melanocyte differentiation in both the NC-derived melanocytes in the embryo and McSC derived melanocytes, but is not required for the specification or establishment of the McSC. We hypothesized that loss of Mitfa activity during zebrafish early development would enrich for the inactive McSC state which could inform us about how McSCs become specified from the NC. Against expectation, we found that loss of Mitfa activity leads to a dramatic expansion of cells at the niche and that these emergent cells line peripheral nerves. These data suggest that Mitfa has previously unknown repressor functions in development that safeguard McSC differentiation programmes.

## Results

### An unexpected repressor function for Mitfa activity at the McSC niche

The *mitfa:GFP* transgene faithfully marks McSCs as they migrate into the niche. Subsequently transgene expression is reduced as McSCs become quiescent (Dooley et al. 2013; Brombin et al. 2022; Brunsdon et al. 2022). We imaged the *mitfa:GFP* transgene in the McSCs in the *mitfa^vc7/vc7^* mutant, which has a temperature sensitive mutation in an intron splice donor site in exon 6 which leads to no Mitfa protein at the restrictive (higher) water temperature and no pigmented melanocytes (referred to as MITF^OFF^ conditions) (**Figure 1B**) (Johnson et al. 2011; Zeng et al. 2015). In these animals, despite the loss of Mitfa protein, the mutant *mitfa* transcript is still expressed, and the *mitfa:GFP* transgene maintains expression (Lister et al. 2014). Our expectation was that, similar to what is seen in wild type zebrafish embryos, McSCs would be found as undifferentiated, single cells in the niche in the absence of Mitfa activity. Against expectation, confocal imaging of zebrafish embryos expressing the *mitfa:GFP* transgene showed dramatically expanded numbers of GFP+ expressing cells at the niche and along the peripheral nerves, forming a chain of GFP+ expressing cells (**Figure 1C**). Notably, unlike in the Mitfa wild type (WT) conditions, whereby the *mitfa:GFP* transgene expression begins to reduce after 48 hours post fertilization (hpf) (Dooley et al. 2013; Brunsdon et al. 2022), we found the GFP+ signal in the absence of Mitfa activity was highly expressed through over three days of development **(Figure 1C)**.

To address if the expansion of GFP+ expressing cells at the niche and along the peripheral nerves was due to the temperature sensitive nature of the *mitfa^vc7/vc7^* mutant allele rather than the loss of Mitfa activity itself, we assessed the expression of the *mitfa:GFP* transgene in the *mitfa^w2^*(*mitfa^nacre^*) allele (Lister et al. 1999), which serves as a second MITF^OFF^ condition due to a loss-of-function point mutation directly in the *mitfa* gene leading to a truncation in the protein. Again, in the absence of Mitfa activity, we detected expansion of the GFP+ signal at the niche and in long chains along nerves (**Figure 1D**).

Thus, in the absence of MITF activity, GFP+ cells in the McSC niche increase in number and expand along nerves dorsally and ventrally in chains that resemble the morphological appearance of Schwann cells (Adameyko et al. 2009; Budi et al. 2011; Dooley et al. 2013; Singh et al. 2016). Together, these data point to a previously unappreciated function for Mitfa in suppressing expansion of McSC progenitors at the niche. While Mitfa activity has well known critical transcriptional activator functions in melanocyte specification directly from the NC across species, we now propose that Mitfa activity includes a repressor function within the McSC lineage to restrain premature expansion of progeny during development.

### scRNA-seq identifies emergent cell populations in MITF^OFF^ embryos

Next, we applied single cell RNA-sequencing (scRNA-seq) to gain insight into the transcriptional states and cellular identity of the cells emerging from the MITF^OFF^ McSCs. We generated *mitfa^vc7/vc7^*embryos that expressed the *crestin:mCherry* neural crest and *mitfa:GFP* melanocyte lineage markers so that we could capture all neural crest and melanocyte lineage cells during establishment of the McSCs. Embryos were maintained in MITF^OFF^ state at 32°C until 24 hpf. We performed droplet-based scRNA-seq on sorted GFP+ and/or mCherry+ cells using the 10x Chromium system (10x Genomics; see Methods) (**Figure 2A**). The embryos used were stage-matched and collected at the same time as those in our previous work allowing for direct comparison to wild type Mitfa+ conditions (MITF^WT^) (Brombin et al. 2022) (**Figure 2B; see Methods**). We analysed the transcriptomes of 1593 MITF^OFF^ untreated cells, 1314 of which passed our quality control (**Supplementary Figure S1A-H**). After clustering via the Louvain algorithm (Butler et al. 2018) and visualisation on a 2D space using the Uniform Manifold Approximation and Projection (UMAP; (McInnes et al. 2018)), we performed cluster calling using a combination of known markers and projections to previously published datasets (Kiselev et al. 2018; Wagner et al. 2018; Saunders et al. 2019; Farnsworth et al. 2020; Brombin et al. 2022) (**Supplementary Figure S1I-K**). We then excluded a further 203 cells that did not seem related to the pigment cell lineage and may be contaminants (namely cells from the hatching gland, muscles and the endothelium) and clustered the remaining 1111 cells as described above (**Figure 2C; Supplementary Figure S1J**).

**Figure 2.**
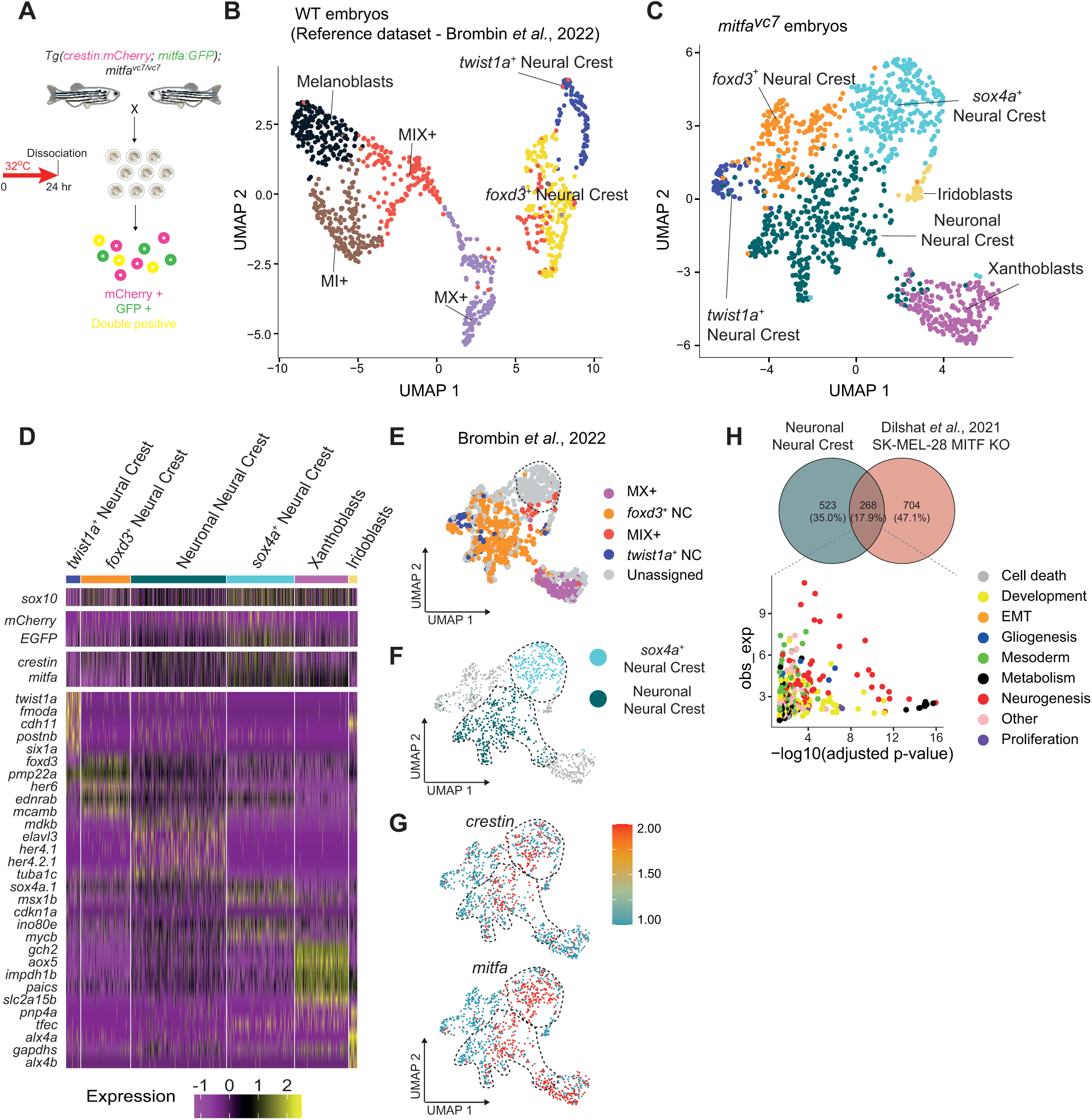
scRNA-seq identifies *de novo* cell populations in MITF^OFF^ embryos. **A.** Schematic of the experimental protocol. Cells that express GFP+, mCherry+ or double GFP+ mCherry+ fluorophores were isolated from *Tg(crestin:mCherry; mitfa:GFP); mitfa^vc7/vc7^* embryos in the MITF^OFF^ state. **B.** UMAP of cells from MITF WT embryos from reference dataset (Brombin et al. 2022). UMAP of GFP+, mCherry+ and double GFP+ mCherry+ cells (n = 996 cells) obtained from zebrafish embryos after Louvain clustering (dims= 20, resolution = 0.5). **C.** UMAP of cells from *mitfa^vc7^* mutant embryos. UMAP of GFP+, mCherry+ and double GFP+ mCherry+ cells (n = 1111 cells) obtained from MITF^OFF^ zebrafish embryos after Louvain clustering (dims= 12, resolution = 0.4). This dataset is stage-matched to *Tg(crestin:mCherry; mitfa:GFP)* WT embryos in B (GEO # GSE178364, (Brombin et al. 2022). **D.** Cell cluster identity. Heatmap showing the average log_2_-fold change expression of five selected genes per cluster identified in C. Log_2_-fold change expression across the 6 clusters of *sox10* and *crestin* (neural crest markers), *mitfa* (melanocyte marker), *mCherry* and *GFP* expression levels are also presented for comparison. **E.** UMAP showing mapping to the MITF^OFF^ Brombin *et al*., 24 hpf dataset. The *sox4a^+^* neural crest cell population demarcated by the dotted line failed to map to any previously identified populations. **F.** UMAP of C labelling the emergent cell populations found in MITF^OFF^ dataset (“*sox4a*^+^ neural crest” and “Neuronal Neural Crest”). **G.** Emergent cell populations maintain neural crest identity. UMAP representations of C with colour change from blue (negative) to red (positive) based on log_2_ mRNA expression of *mitfa* and *crestin* within the “*sox4a^+^* Neural Crest” and “Neuronal Neural Crest” clusters (black dashed line). **H.** Neuronal NC cluster shares a transcriptional profile with MITF KO melanoma cells. Venn’s diagram (top) depicts shared genes between cluster markers of Neuronal NC cell cluster and genes upregulated in SK-MEL-28 MITF KO cell line. Gene ontology (GO) over-representation analysis of shared genes reveals these are related to general development, gliogenesis and neurogenesis (bottom). GO groups were manually grouped into categories (Supplementary Table S4).

In the context of the MITF^OFF^, we identified two neural crest cell (NCC) populations expressing classical NC markers (*e.g. sox10*, *crestin*) and resembling the *twist1a+* NCC and *foxd3+* NCC clusters previously found in the WT dataset ((Brombin et al. 2022), **Figure 2B-E; Supplementary Figure S1; Table S1-S3**). In addition, we identified clusters of cells that expressed transcriptomes consistent with xanthoblasts and iridoblasts (**Figure 2C, D**). Both pigment cell types can originate from *mitfa*-expressing progenitors within the neural crest (Petratou et al. 2018; Dawes and Kelsh 2021; Kelsh et al. 2021; Petratou et al. 2021; Brombin et al. 2022; Brunsdon et al. 2022; Subkhankulova et al. 2023; Brombin and Patton 2024). This is also consistent with an increase in differentiated iridophores in *mitfa^nacre^* embryos at later stages of development (Kelsh et al. 1996; Lister et al. 1999).

Of interest, we found two nascent populations that emerged in the MITF^OFF^ dataset. One cluster we termed “Neuronal NC” as they were closely related to *foxd3+* NC cells in WT embryos but are further enriched for markers for gliogenesis and neurogenesis (*e.g. elavl3, her4.1*). We propose that these cells are an accumulation of undifferentiated cells that would otherwise have become embryonic melanoblasts. A second cluster we termed “*sox4a^+^* NC” because they expressed the stem cell related factor *sox4a* (**Figure 2D-F**). The *sox4a^+^* NC cell cluster comprises cells that predominantly express *mitfa* (**Figure 2G**), suggesting that some of those cells might represent the expanded clusters around the McSC niche. Interestingly, unlike the Neuronal NC cells, the *sox4a*^+^ NC cell cluster could not be mapped consistently to any of the previously identified WT cell populations and appeared to be entirely novel **(Figure 2G; Supplementary Figure S1)**.

We had previously observed neuronal cell states emerging in our zebrafish models of melanoma with low levels of Mitfa activity, and increased *sox4a* expression in melanoma persister cells that lack all Mitfa protein and activity (Travnickova et al. 2019; Travnickova and Patton 2021; Travnickova et al. 2022). This suggested to us that these cell states may be relevant to melanoma disease. To gain insight into the biology of the two emergent cell populations in the MITF^OFF^ dataset, we decided to compare their transcriptomes with an MITF^KO^ SK-MEL-28 human melanoma cell line with a Δ*MITF-X6* genetic deletion, which exhibits a complete loss of MITF protein (Dilshat et al. 2021). We discovered that the Neuronal NC cells shared 268 pathways with MITF^KO^ SK-MEL-28 cells, with many of the overlapping pathways involved in gliogenesis and neurogenesis (**Figure 2H; Table S4**).

Next, we investigated whether *SOX4* could be a direct target of MITF-mediated repression through the analysis of published datasets by Chang and colleagues (Chang et al. 2025). In MITF-high SKMEL28 melanoma cells, genome browser tracks at the *SOX4* locus show that MITF binds multiple regulatory regions and is associated with reduced chromatin accessibility (ATAC-seq), enhancer activity (anti-H3K27ac; CUT&RUN), and mRNA expression (RNA-seq), consistent with transcriptional repression. Loss of MITF (Δ*MITF-X6*) leads to increased accessibility and enhancer activation at MITF-bound sites, accompanied by increased *SOX4* expression, indicating that MITF directly represses chromatin at these loci. Additional regions gain accessibility in Δ*MITF-X6* cells despite lacking detectable MITF binding in wild-type cells, suggesting indirect repression, such as competitive binding by MITF paralogs (Chang et al. 2025; Dias et al. 2025). Consistent with these findings, MITF-low A375 cells display activation of enhancers corresponding to both directly and indirectly repressed regions (anti-H3K27ac; CUT&RUN), similar to the chromatin changes observed following MITF loss in Δ*MITF-X6* cells. Further analysis of RNA-sequencing datasets from A375P cells showed that overexpressing *MITF* resulted in decreased SOX4 expression (Log2FC = –1.97, p = 5.70E-7), while siRNA-mediated knockdown of *MITF* in SK-MEL-28 cells led to increased *SOX4* expression (Log2FC = +0.75, p = 1.37E-5), consistent with a repressor role for MITF at the *SOX4* locus (Dilshat et al. 2021). Thus, human MITF acts as a repressor of *SOX4* in melanoma cells. This regulation is likely context-dependent and influenced by cell differentiation states. We propose that such context-specific repression may reflect conserved developmental pathways governing melanocyte stem cell fate decisions.

### Sox4-positive NC cells form a distinctive node in the developmental trajectory of the NC lineage

To understand lineage relationships between the cell clusters and place the emergent populations within the developmental continuum of the MITF^OFF^ NC lineage, we performed pseudotime analysis which is possible due to the patterned progression of anterior to posterior development and known pigment lineage cell of origin in zebrafish embryos (Brombin et al. 2022) (**Figure 3**; left panel). When we assigned identities to the resulting pseudotime trajectory plot (right panel), we found that the Neuronal NC cells formed distinctive nodes (1, 4) but remained part of the trajectory continuum from early NC cells to differentiation of xanthoblasts. This is consistent with our suggestion that these cells might otherwise normally contribute to the development of the embryonic pattern, and here are likely undifferentiated embryonic melanoblasts. In contrast, *sox4a^+^* NC cells were at the centre of the trajectory scatterplot indicating that those cells are intermediate progenitors, with a singular node (2) marked by a subpopulation of the *sox4a^+^* NC that formed a separate branch in the lineage continuum. We propose these cells are the unexpected *mitfa:GFP+* expressing cell clusters and chains associated with the McSC niche in MITF^OFF^ conditions, as seen in **Figure 1**. We also noticed a portion of *sox4a^+^* NC cells located on the main developmental trajectory (and therefore likely part of normal developmental processes) that is closely related to the xanthoblast population (Mahalwar et al. 2014; McMenamin et al. 2014).

**Figure 3.**
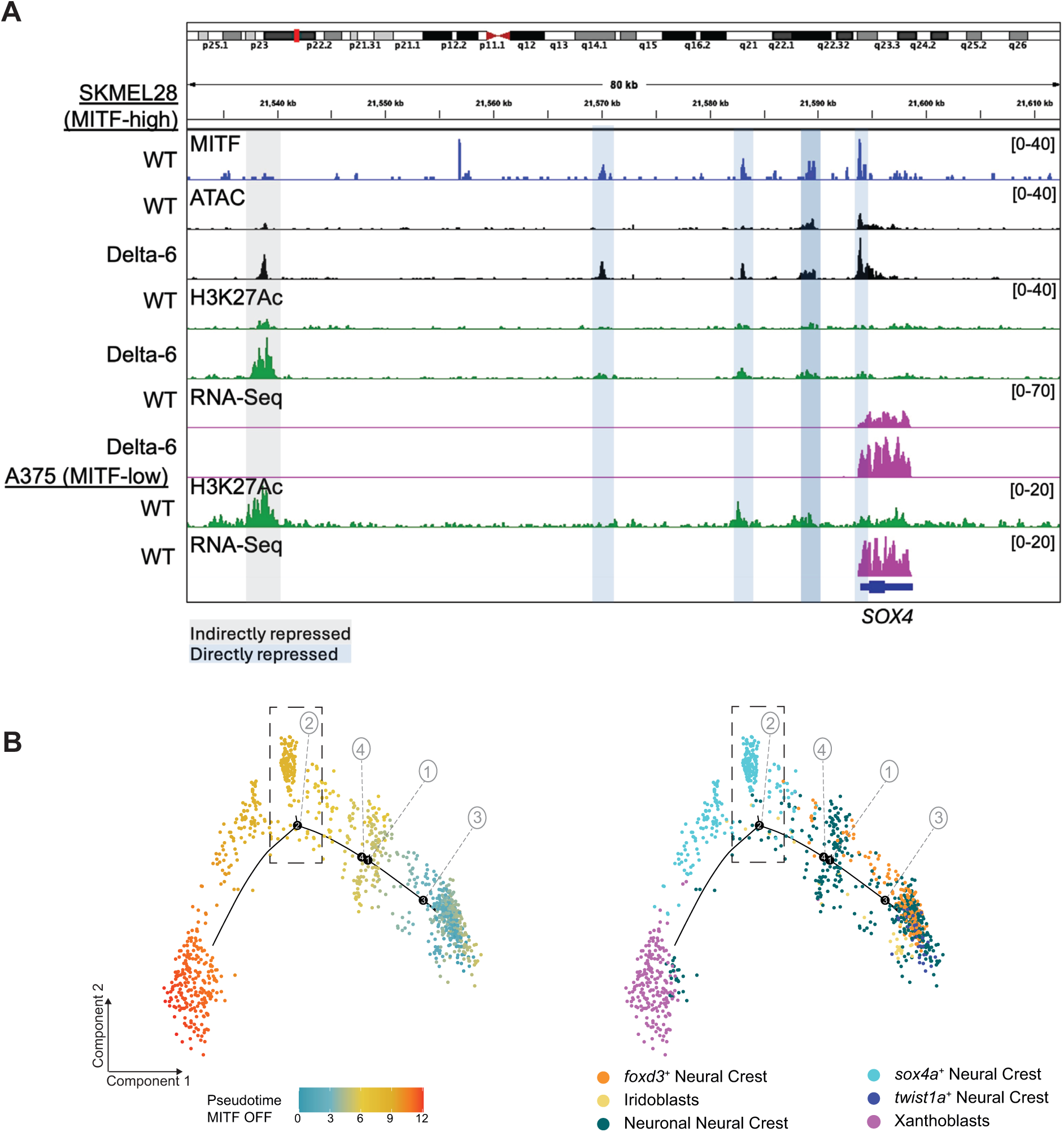
MITF represses chromatin accessibility at the SOX4 locus. **A.** Integrative Genomics Viewer (IGV) tracks at the SOX4 locus in MITF-high SKMEL28 melanoma cells showing MITF binding (anti-MITF CUT&RUN), chromatin accessibility (ATAC-seq), enhancer activity (H3K27ac), and transcription (RNA-seq) in wild-type and MITF-knockout (Δ*MITF-X6*) cells. In wild-type cells, MITF occupies multiple sites at the SOX4 locus and is associated with reduced chromatin accessibility and H3K27ac, consistent with transcriptional repression. Loss of MITF increases accessibility and enhancer activation at MITF-bound sites, indicating that MITF directly represses chromatin at these loci (blue highlight). Additional regions gain accessibility in MITF-knockout cells despite lacking detectable MITF binding in wild-type cells; these are classified as indirectly repressed sites (grey highlight). MITF-low A375 cells similarly show activation of enhancers corresponding to both directly and indirectly repressed regions, consistent with the chromatin changes observed following MITF loss in Δ*MITF-X6* cells. **B.** *Sox4a+* NC cells form a distinctive branch point in the developmental trajectory. Pseudotime ordering of NC and pigment cell lineages of the cells from the MITF^OFF^ dataset (shown in Figure 2C). **Left panel**: Cells are coloured according to their pseudotime scores (from earliest pseudotime point in blue to later pseudotime point in red). **Right panel**: Cells are coloured according to their cluster identity. A subset of the *sox4a^+^* NC cells group in a separate branch (node 2, black dashed rectangle). We propose these to be the axon-associated progenitors in MITF^OFF^ embryos as seen in Figure 1C.

### Mitfa promotes a melanocytic fate while simultaneously repressing alternative fates

We then decided to compare the MITF^OFF^ dataset with its MITF^WT^ counterpart. For this, we integrated the two datasets using the Reciprocal Principal Component Analysis (RPCA) algorithm (Hao et al. 2024) (**Figure 4A**). The two datasets broadly overlapped with cells from both genotypes contributing to most clusters **(Figure 4B, C**). As expected, there was an almost perfect overlap between the early NC cell populations as well as between the MX+ and xanthoblast populations. Conversely, MITF^OFF^ embryos contributed almost exclusively to the iridoblast population – consistent with the increase in iridophores in *mitfa* zebrafish embryonic mutants (Kelsh et al. 1996; Lister et al. 1999) – and to the NC populations expressing neurogenesis markers.

**Figure 4.**
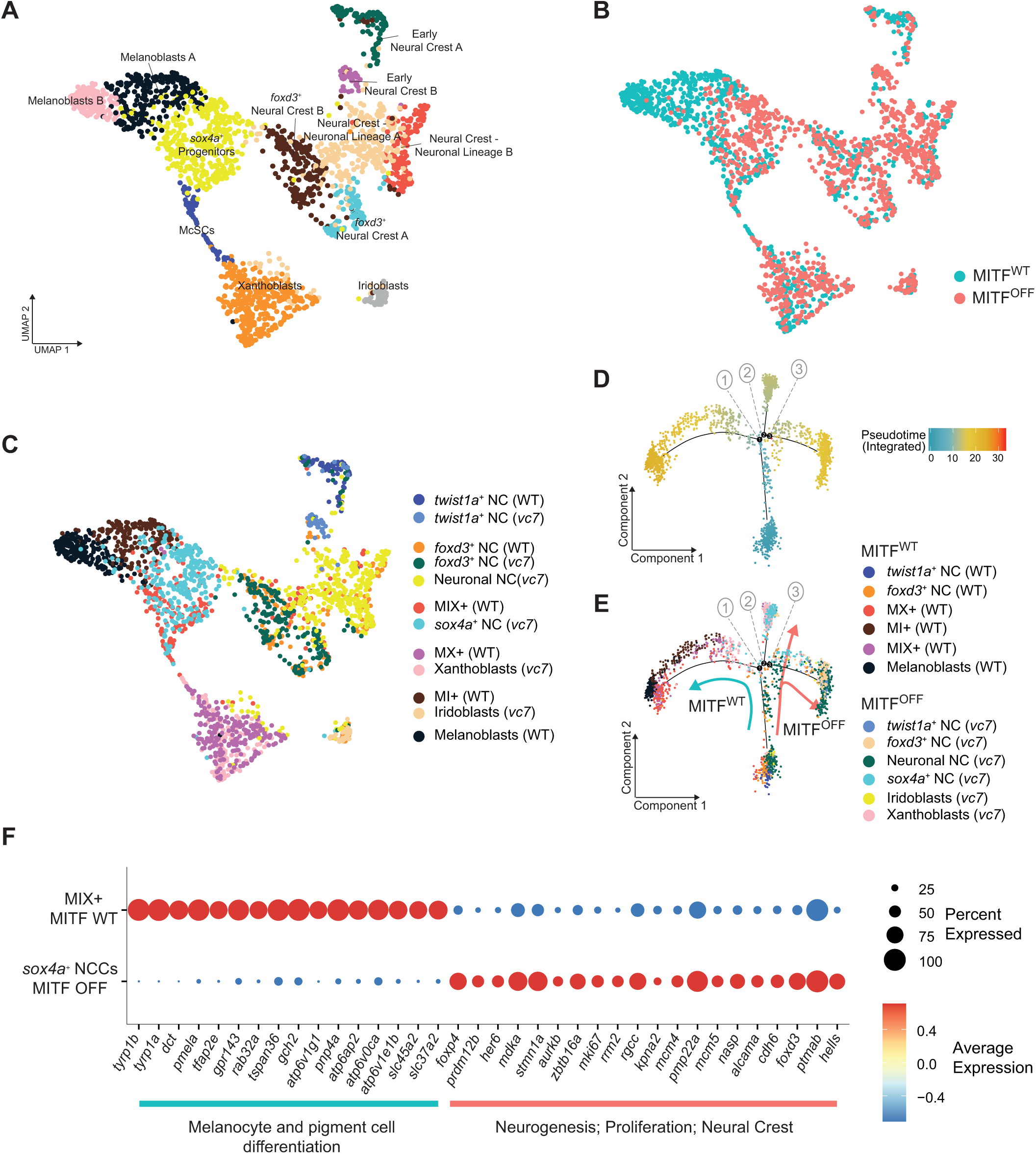
MITF promotes a melanocytic fate while simultaneously repressing alternative fates in the neural crest. **A.** Integration of the wild type and *mitfa* mutant data sets. UMAP of GFP^+^, mCherry^+^ and double GFP^+^ mCherry+ cells from *Tg(crestin:mCherry; mitfa:GFP)* (Brombin et al. 2022) (GEO # GSE178364, 996 cells) and *Tg(crestin:mCherry; mitfa:GFP);mitfa^vc7/vc7^* (1111 cells) obtained from 24 hpf zebrafish embryos after Louvain clustering (dims= 30, resolution = 0.7). Integration was performed following the Reciprocal Principal Component Analysis (RPCA) algorithm within the Seurat (v5.1.0) package (k.anchor = 12). **B.** Integrated dataset by genotype. UMAP representation of the integrated datasets (A) pseudocoloured by genotype. **C.** Integrated dataset by cluster names. UMAP representation of integrated datasets (A) pseudocoloured by original cluster names. **D.** Integrated dataset pseudotime analysis. Combined pseudotime ordering of cells from (Brombin et al. 2022) (GEO# GSE178364, Figure 2B) and the *mitfa^vc7/vc7^* embryos (Figure 2C). The pseudotime was calculated using MITF^WT^ cells, and the most over-dispersed genes were used to order the combined datasets to highlight differences in the melanocyte lineage. Cells are coloured according to their pseudotime scores (from blue to red). **E.** Divergent cell lineage trajectories between genotypes. Pseudotime analysis in D with cells pseudo-coloured according to their cluster of origin. MITF WT trajectory (blue) is distinctive from the MITF^OFF^ trajectory (red). The *sox4a^+^*NC and xanthoblasts in the MITF^OFF^ group in a separate branch at node 2. **F.** McSCs and *sox4a^+^* NC cells have distinct transcriptional programmes. Dotplot showing the log_2_ fold expression of the top genes (grouped by known function) after differential expression analysis (DeSeq2 with ZinbWave correction) between the MIX+ pigment progenitors from the WT dataset (Brombin et al. 2022) GEO# GSE178364, Figure 2B) and the novel cell population (*sox4a^+^* neural crest) identified in the current dataset (Figure 2C). Dot size represents the percentage of cells within cluster expressing the gene and colour indicates average log_2_ fold change expression.

To understand how MITF activity regulates the emergence of pigment precursor cells from the neural crest, we performed an integrated pseudotime analysis between cells obtained from MITF^WT^ and MITF^OFF^ embryos (**Figure 4D, E**). For this, we first constructed developmental trajectory with the cells from the MITF^WT^ embryo only resulting in a “standard” lineage tree as shown in (Brombin et al. 2022) with cells distributed along a pseudotime continuum from the early NCCs to the pigment lineage. The genes that showed the greatest differential expression within clusters were then used to reconstruct the trajectory for the integrated dataset containing both MITF^WT^ and MITF^OFF^ cells thereby allowing for direct comparison between genotypes. Early NCCs (*foxd3^+^* and *twist1a^+^* NCCs) from both genotypes grouped together on the root of the integrated pseudotime trajectory indicating that MITF activity doesn’t affect early stages of neural crest specification. Pseudotime ordering then markedly differed at the pigment progenitor state (node 1, **Figure 4E**) with all the remaining cells from MITF^WT^ embryos grouped in a single branch of the pseudotime trajectory leading to differentiated melanocytes, while most of the cells from MITF^OFF^ embryos accumulated in separate branches (node 2 and node 3) as undifferentiated cells. Notably, most of the *sox4a^+^*NCCs grouped in the branch starting from node 2, reinforcing the distinct nature of these cells. Xanthoblast progenitors also accumulated here likely supporting the fact that MITF activity is required for the specification of the xanthophores in zebrafish (Miyadai et al. 2023).

Interestingly, some of the MITF^OFF^ *sox4a+* NC population overlaps with MITF^WT^ MIX+ cells which includes McSCs **(Figure 4C)**. To gain greater insight into these common cell identities, we performed differential expression analysis between MITF^OFF^ *sox4a+* NC population and the MITF^WT^ MIX+ cells. Despite sharing some overlap in the 2D UMAP space, the MITF^OFF^ *sox4a+* NC transcriptomic landscape differed from MITF^WT^ MIX+ cells expressing neurogenesis, proliferation and neural crest markers while the MITF^WT^ MIX+ cells expressed MITF targets, such as melanin synthesis related *dct* and *tyrp1a/b* (**Figure 4F; Table S5**). These data suggests that loss of MITF activity can actively shape a new transcriptional identity of the McSCs and their progenitors.

### Lineage tracing the embryonic McSC through juvenile and early adult stages

Next, we wanted to understand how cells from the McSC contributed to the adult pigment pattern in the absence of MITF activity. To this end, we designed an experiment to study the impact of MITF activity on the McSC lineage by coupling lineage tracing with imaging and scRNA-seq (**Figure 5A**). We first tried the *mitfa* promoter upstream of a tamoxifen-inducible *creERT2* element (*mitfa: creERT2*); however, we mostly labelled the embryonic developing melanocytes and were unable to identify a developmental time point where we could specifically label the McSCs, likely due to low transgenic *mitfa* expression levels in the McSC (*data not shown*).

**Figure 5.**
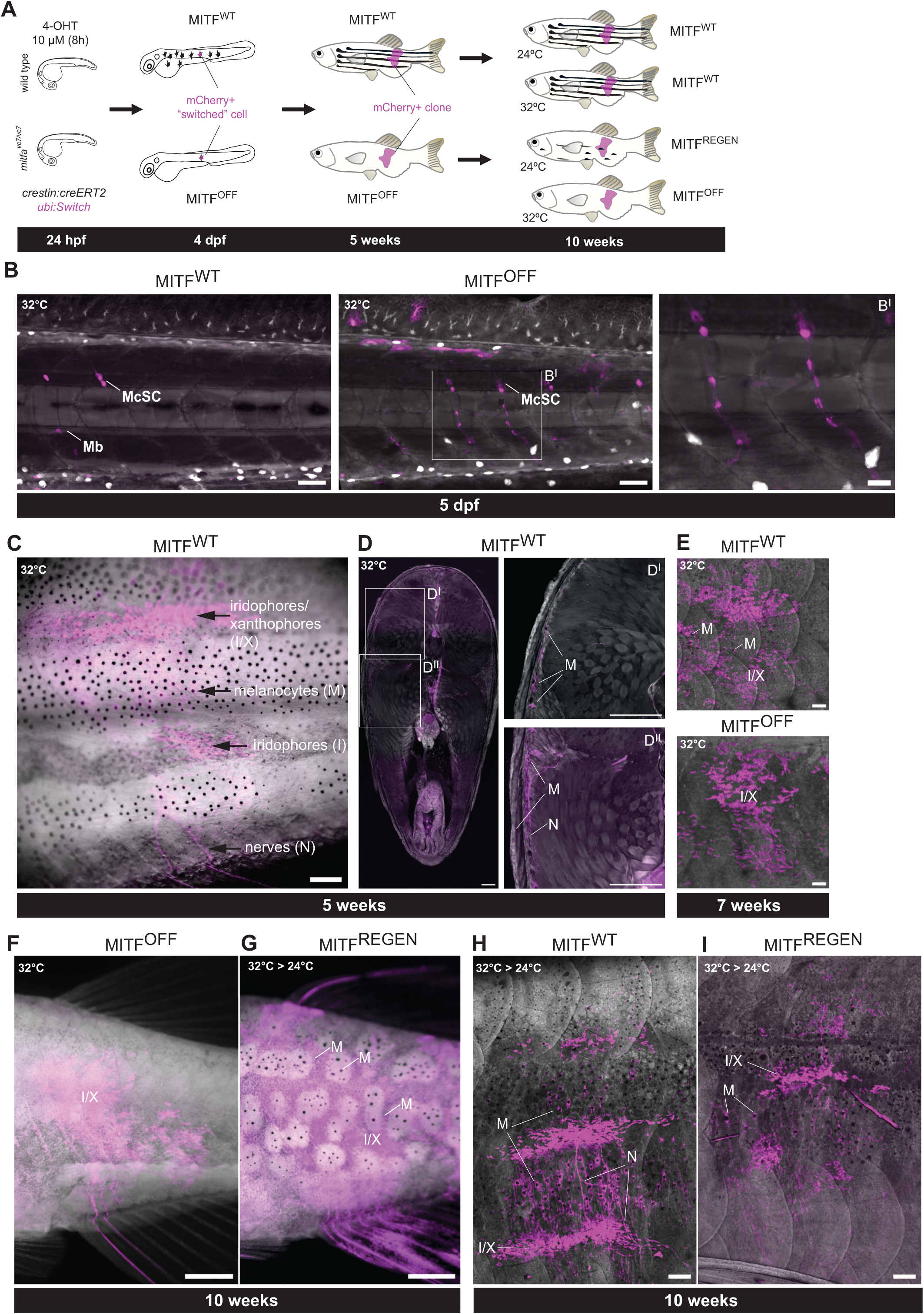
Tracing McSCs into the adult pigment pattern and altering MITF activity. **A.** Overview of experimental protocol. Experimental protocol for *crestin:creERT2* mosaic lineage tracing of McSCs derivatives with or without MITF activity, and in regeneration. *Tg(crestin:creERT2; ubi:Switch)* in wildtype and *mitfa^vc7/vc7^* were raised until 24 hpf at the non-permissive temperature 32°C before being treated with 10 µM 4-Hydroxytamoxifen (4-OHT) for 8 hours. Treated embryos of both genotypes were raised at 32°C for 5 weeks to generate adult fish with MITF^WT^ or MITF^OFF^ lineage traced clones. In addition, the 5 week old WT and *mitfa^vc7/vc7^* fish were raised for a further 5 weeks at either 32°C (MITF^OFF^) or shifted to 24°C to restore Mitfa activity (MITF^REGEN^). Only treated embryos displaying mCherry+ McSCs were raised beyond 5 dpf. mCherry+ is shown in magenta in the schematic. **B.** *Crestin:creERT2* mosaic lineage tracing of McSCs derivatives with or without MITF activity during zebrafish development. McSCs are indicated, and the chains of McSC progeny along nerves are visible in the MITF OFF conditions (zoomed image; right). Within our experimental conditions, *crestin:creERT2* marks a few other cells at this stage including the occasional embryonic derived melanoblast (Mb). White pseudo-colouring is used for the GFP channel and magenta for the mCherry channel. Scale bars: 50 µm. (I) Scale bar: 20 µm. **C.** *Crestin:creERT2* mosaic lineage tracing of McSCs into the adult pigment pattern (5 weeks). A clone of all three pigment cell types and nerve cells that have been traced from a McSC in the embryo. Scale bars: 200µm. **D.** Confocal stacks of vibratome transversal sections obtained from fish at the stages shown in C. mCherry^+^ cells are detectable both around the nerves (N, zoomed image D’’) and in pigmented melanocytes (M, zoomed image D’) at the skin and scales. Scale bars: 200 µm. **E.** Images of MITF^WT^ and MITF^OFF^ skin at 7 weeks. Melanocytes are clearly visible in the MITF^WT^ skin, while no melanocytes are seen in the MITF^OFF^ skin. Scale bars: 100 µm. **F.** No melanocytes are present in *mitfa^vc7/vc7^* fish raised at the non-permissive temperature (MITF^OFF^) at 10 weeks. Scale bar: 1 mm **G.** Melanocytes in regenerating *mitfa^vc7/vc7^* (kept at 32°C for 5 weeks and then downshifted at 24°C for other 5 weeks) (MITF^REGEN^). Melanocytes emerge in clusters that will coalesce into stripes emerge. Representative images of 7-8 animals imaged per condition. Scale bar: 1 mm. **H.** Melanocytes, iridophores, xanthophores and nerve associated cells in MITF^WT^ skin at 10 weeks (kept at 32°C for 5 weeks and then downshifted at 24°C for other 5 weeks). Scale bar: 200 µm. **I.** Melanocytes in regenerating *mitfa^vc7/vc7^* at 10 weeks (kept at 32°C for 5 weeks and then downshifted at 24°C for other 5 weeks) (MITF^REGEN^). Scale bar: 200 µm.

Next, we considered *crestin:creERT2* because our imaging with the *crestin:mCherry* construct showed sustained expression in McSCs while its expression faded in other cells at 24 hpf (Brombin et al. 2022; Brunsdon et al. 2022). We cloned the promoter of the *crestin* gene upstream of a tamoxifen-inducible *creERT2* element (*crestin:creERT2*) and injected it into the *ubi:Switch* transgenic line in either the WT or *mitfa^vc7/vc7^*background (Mosimann et al. 2011; Kaufman et al. 2016; Brombin et al. 2022; Brunsdon et al. 2022). *crestin* is expressed early in development and labels most of the migrating trunk neural crest (Luo et al. 2001)(Kaufman et al. 2016). Thus, to achieve recombination within the McSCs, we decided to start the 4-OHT induction at 24 hpf, just after McSC establishment at the stem cell niche (Brombin et al. 2022) thereby uncoupling the McSC lineage from the embryonic neural crest. Further, we used a lower dose of 4-OHT (10 µM for 8 hours) **(Figure 5A)** to allow for partial (mosaic) recombination within the McSC population and thereby facilitate lineage tracing of the “switched” McSCs (GFP+ to mCherry+). Recombined McSCs (mCherry^+^) could be detected at the niche at 4 dpf in both MITF^WT^ and MITF^OFF^ embryos **(Figure 5B)**. As we had seen with the *mitfa:GFP* transgene, the *crestin:creERT2* switched McSCs had expanded progenitors in the MITF^OFF^ state.

Due to the treatment of 4-OHT at 24 hpf, when *crestin* expression is declining, mCherry+ could be detected in only a limited number of other cell types in both genotypes. Therefore, we selected for embryos that had switched McSCs, and grew the fish at 32°C for 5 weeks until the MITF^WT^ juveniles reached the end of metamorphosis (*i.e.* they displayed clearly visible stripes). We identified mCherry+ MIX+ clones (containing <u>m</u>elanocytes, iridophores and <u>x</u>anthophores and nerve associated cells) on the flank of metamorphic MITF^WT^ fish **(Figure 5C)**. This indicates that the *crestin* transgene faithfully labels the multipotent adult McSC in the embryo, and that these cells contribute to clones of pigment cells and nerve associated cells in the adult.

We then performed serial transversal sections of lineage traced fish at the site of the mCherry+ clone. mCherry+ cells were found in association with nerves as well as in differentiated pigment cells (**Figure 5D**). In contrast, in the MITF^OFF^ conditions, we found IX clones on the flank but not melanocytes (**Figure 5E**). Thus, loss of MITF activity does not prevent the development of McSC derivatives other than melanocytes in the adult.

### MITF activity restores melanocytes in zebrafish after long term MITF depletion

We then revisited our lineage tracing experiment in the context of regenerating melanocytes, called MITF^REGEN^, where we restored MITF activity from OFF to ON states (**Figure 5A**). At 5 weeks, fish were divided into two batches per genotype and maintained for a further 5 weeks either at the non-permissive temperature (32°C) (**Figure 5F**) or shifted to the permissive temperature (24°C) where regenerating melanocytes begin to coalesce into stripes **(Figure 5G)**. We analysed the mCherry+ clone composition on the skin of the fish by confocal imaging and found that the clones in MITF^REGEN^ fish contained newly developed melanocytes, similar to their wild type counterparts (**Figure 5H, I**). This finding indicates the McSC lineages retain the potential to generate melanocytes even after long-term MITF inactivity.

### ScRNA-seq of lineage-traced fish reveals multiple populations of McSC lineage progenitors

To evaluate the impact of MITF activity on the McSCs and its derivatives, we performed scRNA-seq on isolated mCherry+ cells from the lineage tracing experiment detailed above that enriches for McSC lineage progenitors. We isolated mCherry+ cells from four vc7 fish under regenerating conditions (MITF^REGEN^), four maintained as controls at 32°C (MITF^OFF^), and combined 8 wild-type fish raised at either 32°C or 24°C to represent MITF^WT^. We focused analysis on the pigment cells from the trunk by removing the head and internal organs and performed cell dissociation of the remaining tissues including muscle, fins and skin. Accurate sorting of mCherry+ cells was experimentally determined using fish lacking transgene and non-treated ubi:Switch fish from both genotypes **(Supplementary Figure S2).** We sequenced the transcriptome of 3380 mCherry+ cells, 3257 of which passed our quality control (**Supplementary Figure S3A-B**). 597 cells were excluded from the downstream analyses after cluster calling (presumed contaminants: blood cells, vessels, unknown ciliated cells). We then re-clustered the remaining 2660 cells as described above and we identified 13 clusters (**Figure 6A**; **Table S6; Supplementary Figure S3C-D**). We then mapped the transcriptomes of the sorted cells to known datasets and used known markers to perform cluster calling (**Figure 6A, B; Table S7; Supplementary Figure S3E**). Five clusters contained cells that are not usually associated with the pigment lineage *per se* (kidney cells, mesenchyme and lateral line). Nevertheless, we decided to retain them because they contain neural crest derivatives. The remaining clusters were in keeping with pigment progenitors within the McSC lineage.

**Figure 6.**
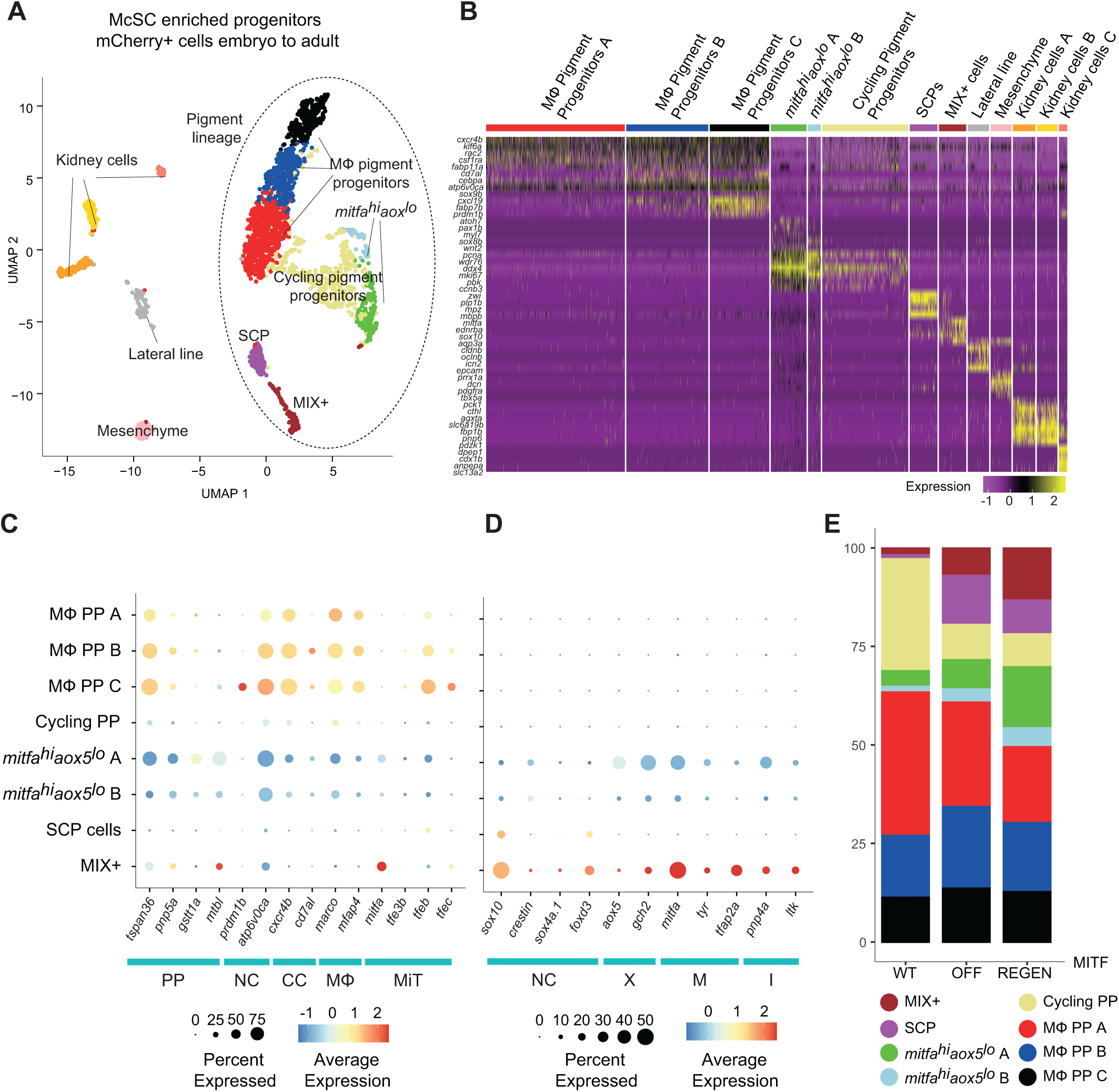
scRNA-seq of McSC derivatives in MITF^ON^, MITF^OFF^ and regenerating fish. **A.** UMAP of mCherry+ cells (n = 2660 cells) obtained from 10 weeks old *Tg(crestin:creERT2; ubi:Switch)* and *Tg(crestin:creERT2; ubi:Switch); mitfa^vc7/vc7^* zebrafish treated with 4-OHT after Louvain clustering (dims= 9, resolution = 0.5). **B.** Heatmap showing the average log_2_-fold change expression of four selected genes per cluster identified in **A**. **C.** Dot plot representing the average expression of “ML Pigment Progenitors” – associated genes across the clusters. CC: chemokine genes, NC: neural crest genes, ML: macrophage markers, MiT: MiT genes, PP: markers expressed in the “Pigment Progenitors” population identified by (Saunders et al. 2019). Dot size indicates the percentage of cells within cluster expressing the gene and colour represents the expression level. **D.** Dot plot representing the average expression of pigment lineage genes across the clusters. I: iridophores, M: melanocytes, NC: neural crest genes, X: xanthophores. Dot size indicates the percentage of cells within cluster expressing the gene and colour represents the expression level. **E.** Bar plot representing the differences in the proportions of cells from the pigment lineage within each cluster from MITF^WT^, MITF^OFF^ and MITF^REGEN^ fish.

Unfortunately, we didn’t detect any mature pigment cells, which may have been lost during our processing of the skin. However, given our transversal sections of lineage traced fish at the site of the mCherry+ clone that showed progenitors within the body tissues **(Figure 5D)**, we continued to explore the McSC lineage progenitors in MITF^WT^, MITF^OFF^ and MITF^REGEN^ conditions. We identified two clusters of pigment progenitors that mapped to the *mitfa^hi^aox5^lo^* precursors that are activated during melanocyte regeneration, described in (Frantz et al. 2023b) (**Supplementary Figure S4A**). As their name suggests, these cells express *mitfa* and low-to-no *aox5* (a xanthophore marker) and are highly proliferative (**Figure 6B; Supplementary Figure S4B**). Next, we identified a populations of actively cycling cells (Cycling pigment progenitors), based on expression of G1/S and G2/M marker genes (Sur et al. 2023)(**Figure 6B; Supplementary Figure S4B**) as well as Schwann Cell Progenitors based on SCP marker expression. Furthermore, we found a population of cells expressing high levels of NC, all pigment cell markers, as well as *sox4*, called “MIX+” (for <u>m</u>elanocytes, iridophores and <u>x</u>anthophores) **(Figures 6B; Supplementary Figure S4A**).

Within the mCherry+ lineage traced cells, we found three intriguing populations whose transcriptomes were very similar and mapped to both the pigment progenitors described by Saunders and colleagues but also to the tissue resident macrophages (TRMs) described by Frantz and colleagues (Saunders et al. 2019; Frantz et al. 2023b) **(Supplementary Figure S4A)**. After excluding the possibility that these cells were doublets by looking at differences in Unique Molecular Identifiers (UMIs) and the variety of expressed genes **(Supplementary Figure S4B)**, we thoroughly investigated their transcriptomes and found that these cell populations expressed both NC and iridophore progenitor markers (*tfec*) as well as markers traditionally associated with macrophages, including notably, *mfap4*, *marco, and tfeb* (but not *mpeg1*), as well as genes associated with macrophage activation and migration (**Figure 6B-D)**. *tfec*, was particularly highly expressed and this might indicate that these cells are most closely related to the iridophore lineage because *tfec* is necessary for the specification of the iridophores from the NC at embryonic stages (Petratou et al. 2018; Petratou et al. 2021; Brombin et al. 2022). We named these macrophage pigment progenitors (MΦ PP) to reflect the names from Saunders and colleagues, and Frantz and colleagues (Saunders et al. 2019; Frantz et al. 2023b)

### Schwann cell progenitors and MIX+ cells are enriched, while Cycling progenitors are reduced in the absence of MITF activity

We examined the cell states that varied between the MITF conditions. The *mitfa^hi^aox5^lo^* cells showed expansion in the MITF^REGEN^ conditions, suggesting that the *mitfa^hi^aox5^lo^* cells are McSC progenitors that contribute to stripe regeneration **(Figure 6E; Supplementary Figure S4C**), possibly as skin-resident progenitors (Chang et al. 2025). In contrast, the cycling cells which made up almost 25% of the pigment progenitors in the MITF^WT^ fish, were dramatically depleted in both the MITF^OFF^ and MITF^REGEN^ conditions, indicating that these cells may be a cycling pool of MITF-dependent melanocyte progenitors in growing fish. The SCP and MIX+ cells were enriched in the MITF^OFF^ and MITF^REGEN^ conditions, suggesting that these cells are progenitors that, in wild type conditions, are slowly depleted over time to produce melanocytes.

## Discussion

How embryonic melanocyte stem cells generate large clones of pigment cells in the adult zebrafish represents a fundamental question in the biology of animal pigmentation and patterning. Here, we find an unexpected role for MITF repressor activity in the McSC lineage in zebrafish embryos, leading to emergent long chains of *mitfa:GFP* expressing cells along nerves, reminiscent of SCPs. Lineage tracing these cells to adulthood demonstrates that without Mitfa these cells can still generate iridophores and xanthophores, as well as nerve associated cells, and a previously undescribed population expressing both iridophore and macrophage markers but are unable to generate differentiated melanocytes. Our zebrafish studies reveal that MITF has a dual function in zebrafish melanocyte developmental biology: as an activator in the differentiating melanocyte lineages and as a repressor in McSCs.

We find that in the absence of MITF activity, there is an accumulation of SCP-like cells in the embryo based on the cell shape, as well as in the adult based on the gene expression. Using *sox10* or *tfap2b* as a lineage tracer, or the *mitfa:GFP* transgenic line, SCP-like cells have been previously shown to emerge from the McSC niche and the marked progenitors to form long chains of cells along nerves to pigment the adult zebrafish (Dooley et al. 2013; Singh et al. 2016; Brombin et al. 2022). Here, we find that a new function for Mitfa to directly repress the expansion of these SCP-like cell populations. We find two emergent transcriptional cell states in the absence of MITF: a neuronal cell state similar to the MITF null cells (Dilshat et al. 2021); and a *sox4^+^*cell state that may have some similarity to McSCs, but with a distinct transcriptional identity. Given the overlap in UMAP space of the McSCs (WT) and *sox4^+^* cells (*mitfa^vc7^*) by scRNA-sequencing as well as the pseduotime analysis, we propose that the *sox4^+^* cells reflect the long chains of cells along nerves that we detect by imaging in the *mitfa^vc7^* mutant zebrafish. In contrast, we propose that the Neuronal NC cell state may represent the embryonic melanocyte lineage directly from the neural crest in the absence of the MITF activity. Analysis of the regulatory regions, chromatin accessibility, enhancer activity and expression of *SOX4* shows that *SOX4* is both a direct and indirect target for MITF in some human melanoma cell lines. Taken together with our data presented here in zebrafish embryos, we hypothesize that the SCP-like cell states are negatively regulated by Mitfa in zebrafish McSCs likely through repression of *sox4*.

These emergent cell states observed in the *mitfa* mutants may reflect the phenotypic state shifts seen in melanoma upon loss of MITF activity. In melanomas from patients, low MITF transcriptional activity is associated with poor outcomes (Cancer Genome Atlas 2015; Lauss et al. 2016), and we have previously shown that MITF-low states in zebrafish melanoma are oncogenic and promote melanoma cell invasive phenotypes coupled with neuronal and invasive gene expression programmes (Travnickova et al. 2019). Recent evidence suggests that the molecular mechanisms underlying loss of MITF at genome binding sites enables competitive binding by MITF paralogs that drive distinct gene expression programmes, including mesenchymal and metastatic cell states (Chang et al. 2025; Dias et al. 2025). It will be of interest to determine if similar antagonistic programmes with MITF paralogs function in the context of developmental biology and McSC lineages.

Through fate mapping from the McSC to the adult, using *crestin:creERT2*, we provide further evidence that single McSCs are multi-potent and generate clones of cells in the adult zebrafish (Dooley et al. 2013; Singh et al. 2016; Brombin et al. 2022). As expected, we find Mitfa is critical for differentiation of the melanocyte lineage. Using scRNA-sequencing, we reveal that heterogeneity from this clone in the adult includes SCP-like cells, MIX+ cells, as well as cycling progenitors and *mitfa^hi^aox5^lo^* cells that have been previously detected in the skin (Frantz et al. 2023a). Our Mitfa depletion and restoration experiments showed that the SCP-like and MIX+ cells accumulate in the absence of Mitfa, while the proliferating melanocyte progenitors decrease, suggesting that SCP-like and MIX+ cells may directly contribute to the cycling progenitor pool. Further, the MIX+ cell population expands during regeneration, along with one of the *mitfa^hi^aox5^lo^*populations, supporting previous studies suggesting that regenerating melanocytes in the adult may depend on local subpopulations (Iyengar et al. 2015; Frantz et al. 2023a). We also identified a cell population that expresses macrophage and iridophore markers, perhaps reflecting an immune-like function for pigment lineage cells that is not dependent on Mitfa, but is part of a shared clonal linage that originates with McSCs.

Lineage tracing of the McSC in the embryo to the adult zebrafish has not previously been interrogated by conditional genetic perturbation, and our study comes with some limitations. We used the neural crest *crestin:creERT2* line, rather than a line that is specifically expressed in the McSC because McSC specific reporter lines are not currently available. While we carefully selected animals where we could only detect expression in the McSCs, other neural crest cells and their lineage may have been included in our analysis. A further limitation is that we did not recover differentiated melanocytes from the skin of our adult zebrafish, which likely reflect cell isolation conditions leading to cell loss during the scRNA-sequencing process.

In summary, we show that Mitfa repressor activity restrains McSCs from premature expansion during development, likely through repression of *sox4a*, and that McSCs retain multi-potency in the absence of Mitfa except for melanocyte differentiation. These observations were made possible by the combination of imaging and single cell technologies that reveal cellular transcriptional states and expansions of McSC progeny along nerves. These findings may have broader implications for the impact of loss of MITF activity in human genetic disease, including Waardenburg Syndrome, Teitz and COMMAND syndromes (OMIM 617306, 103500, 193510), as well as in melanomas with low to no MITF protein expression, because despite the absence of MITF activity, the *MITF*-expressing cell lineages may expand and generate new cell states that contribute to disease. In addition, within the zebrafish community, studies that use *mitfa* mutants (*e.g. nacre*) (Lister et al. 1999; Zeng et al. 2015) may consider that in addition to loss of differentiated melanocytes, the animals have emergent cell states. In conclusion, our study provides the first insight into a new developmental phenotype that emerges in the absence of Mitfa function, reflecting an unexpected repressor function for Mitfa in the melanocyte stem cell lineage.

## Author Contributions

Conceptualisation: AB, TC, EEP; Methodology: AB, TC, CW, SM, JT, HB, ER, HNV, CK; Software: AB, JT, HB; Validation: AB; Formal analysis: AB, JT, HNV, CK Investigation: AB, CW, SM; Resources: ES, CK, EEP; Writing original draft: AB, EEP; Writing review and editing: All authors; Visualisation: AB; Supervision: ES, CK, TC, EEP; Funding acquisition: TC, CK, ES, EEP.

## Acknowledgements

We are grateful to Colin Goding (University of Oxford) for helpful discussions, and the University of Edinburgh Institute of Genetics and Cancer Zebrafish Facility, Flow Cytometry Facility and Imaging Facility. ES is funded by the Icelandic Research Fund, [239941] and HNV by a postdoctoral grant [2511591]. CK is funded by the Melanoma Research Alliance [1426690] and the National Cancer Institute [R03-CA297549]. EEP acknowledges support from the Medical Research Council [MC_UU_00035/13], the Melanoma Research Alliance and Rosetrees Trust [MRA Awards 917226, MYIA\100003], the Wellcome Trust [333845/Z/25/Z] and by the Cancer Research UK Scotland Centre [CTRQQR-2021\100006].

---

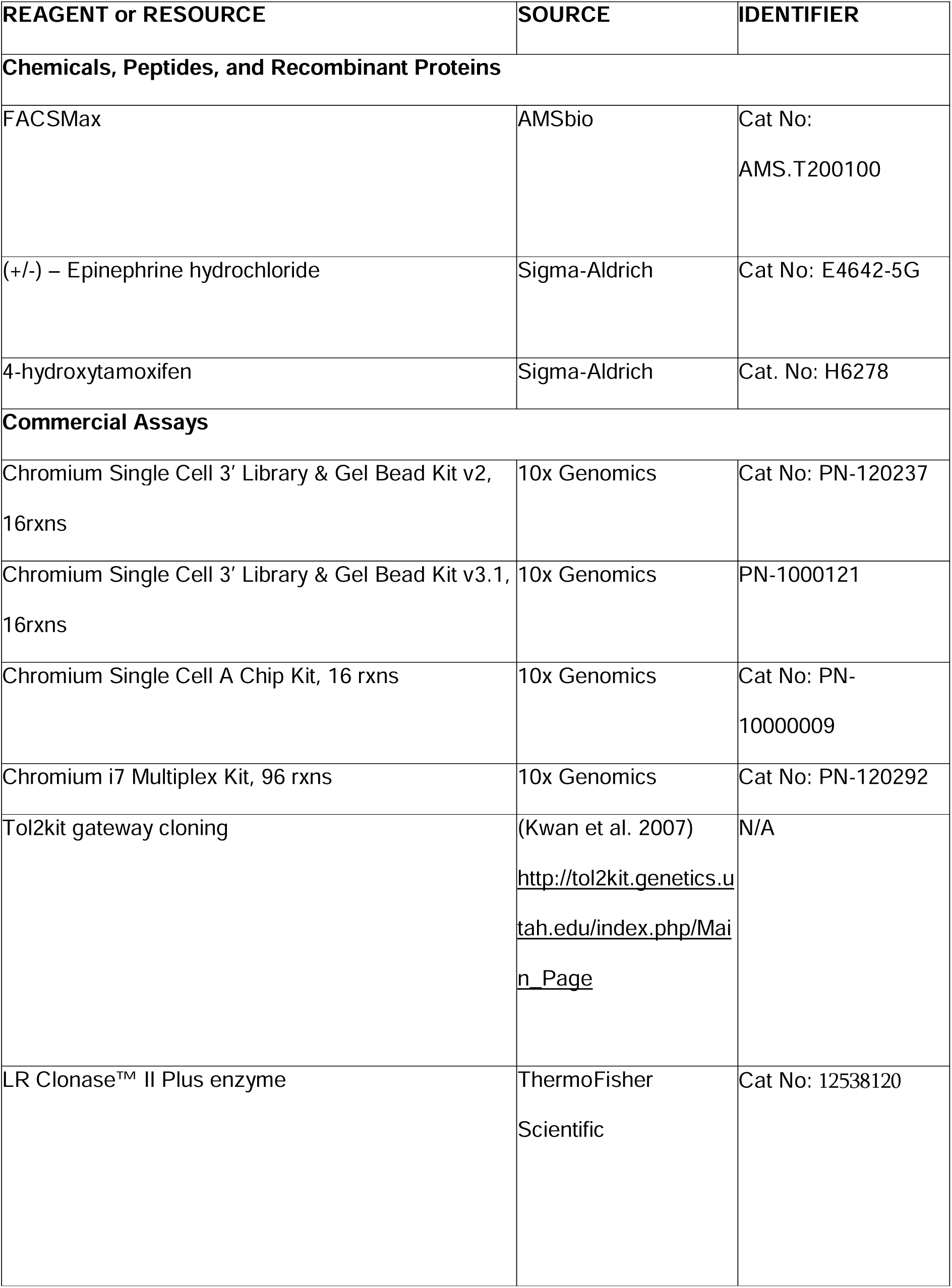
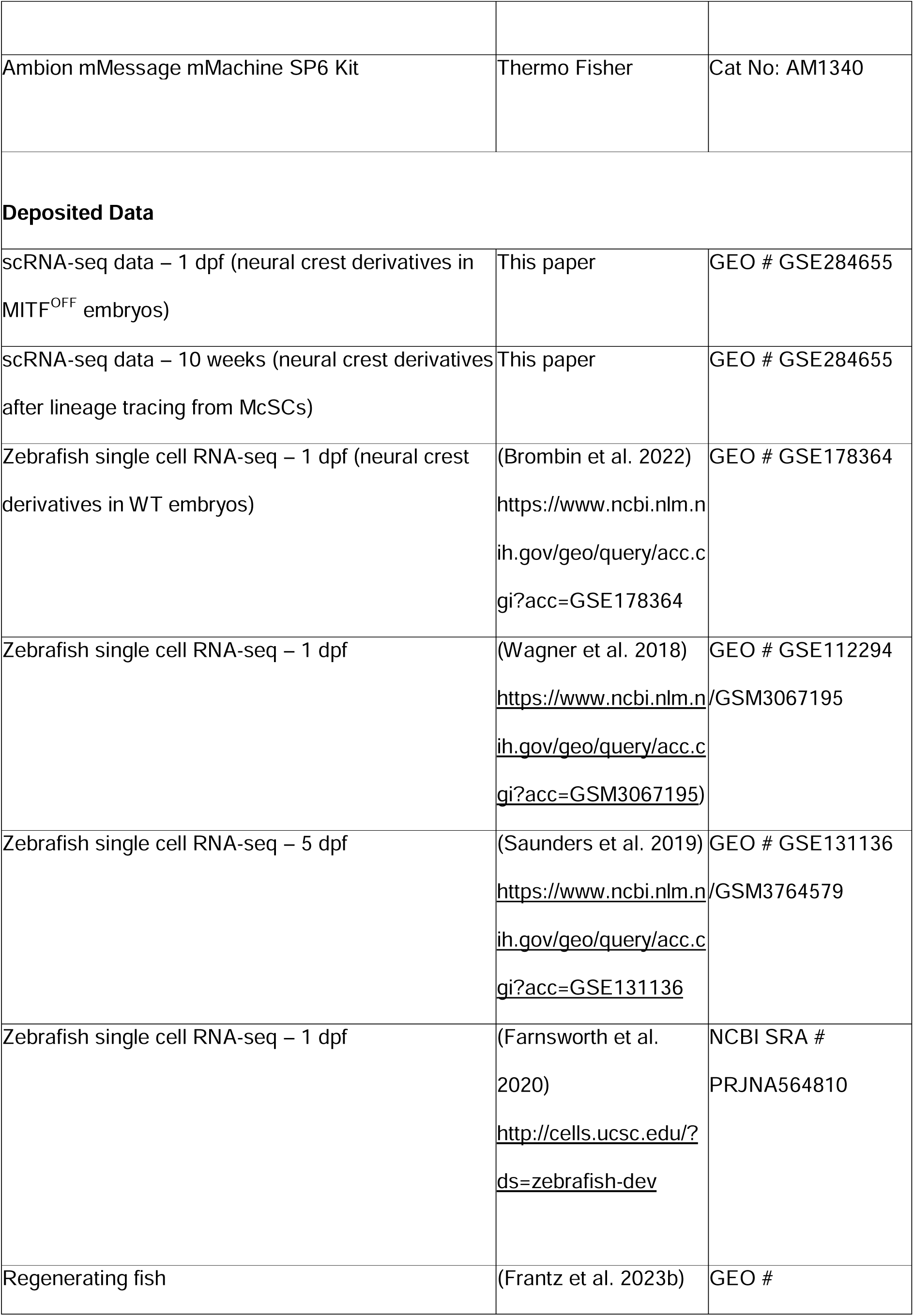
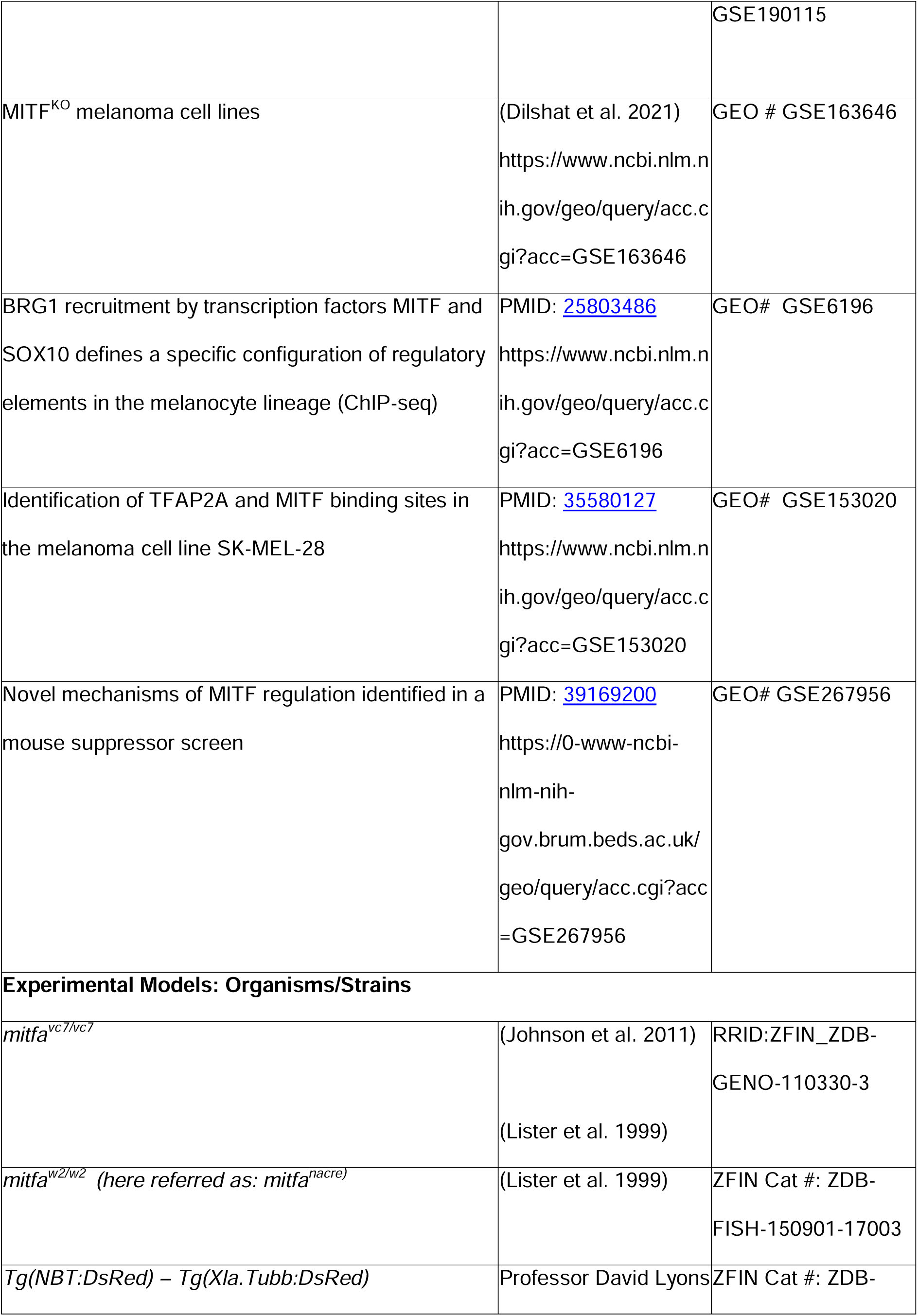
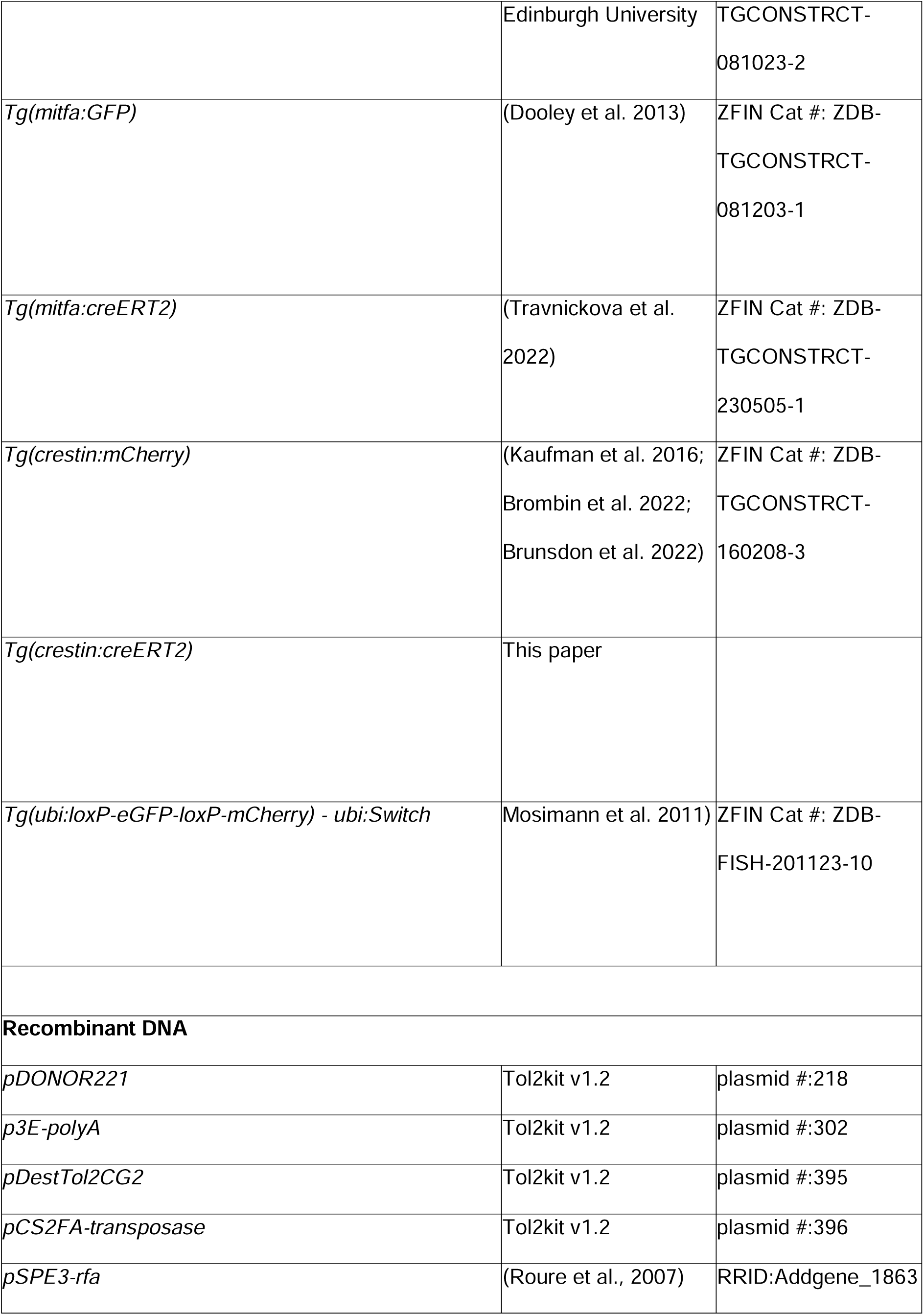
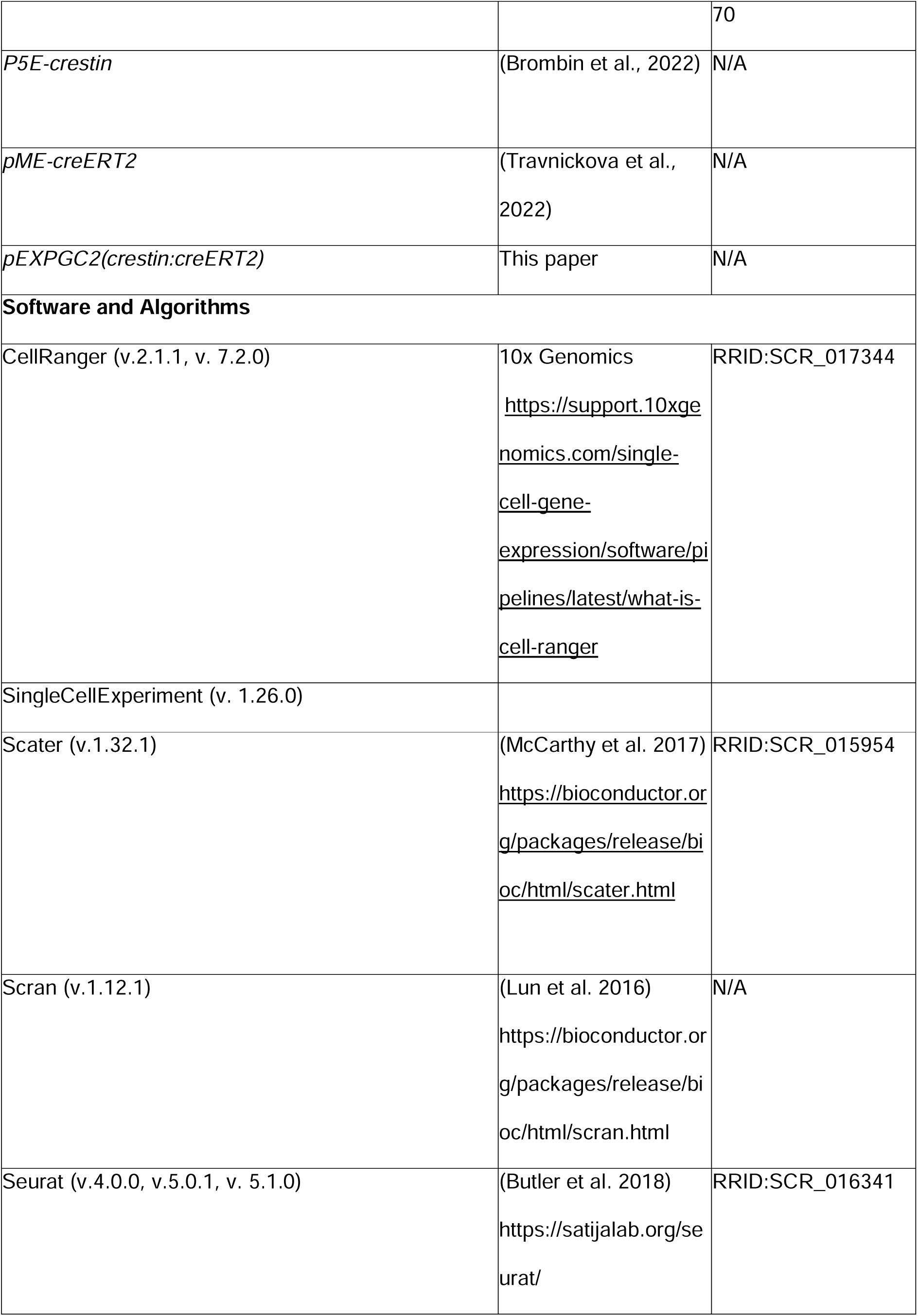
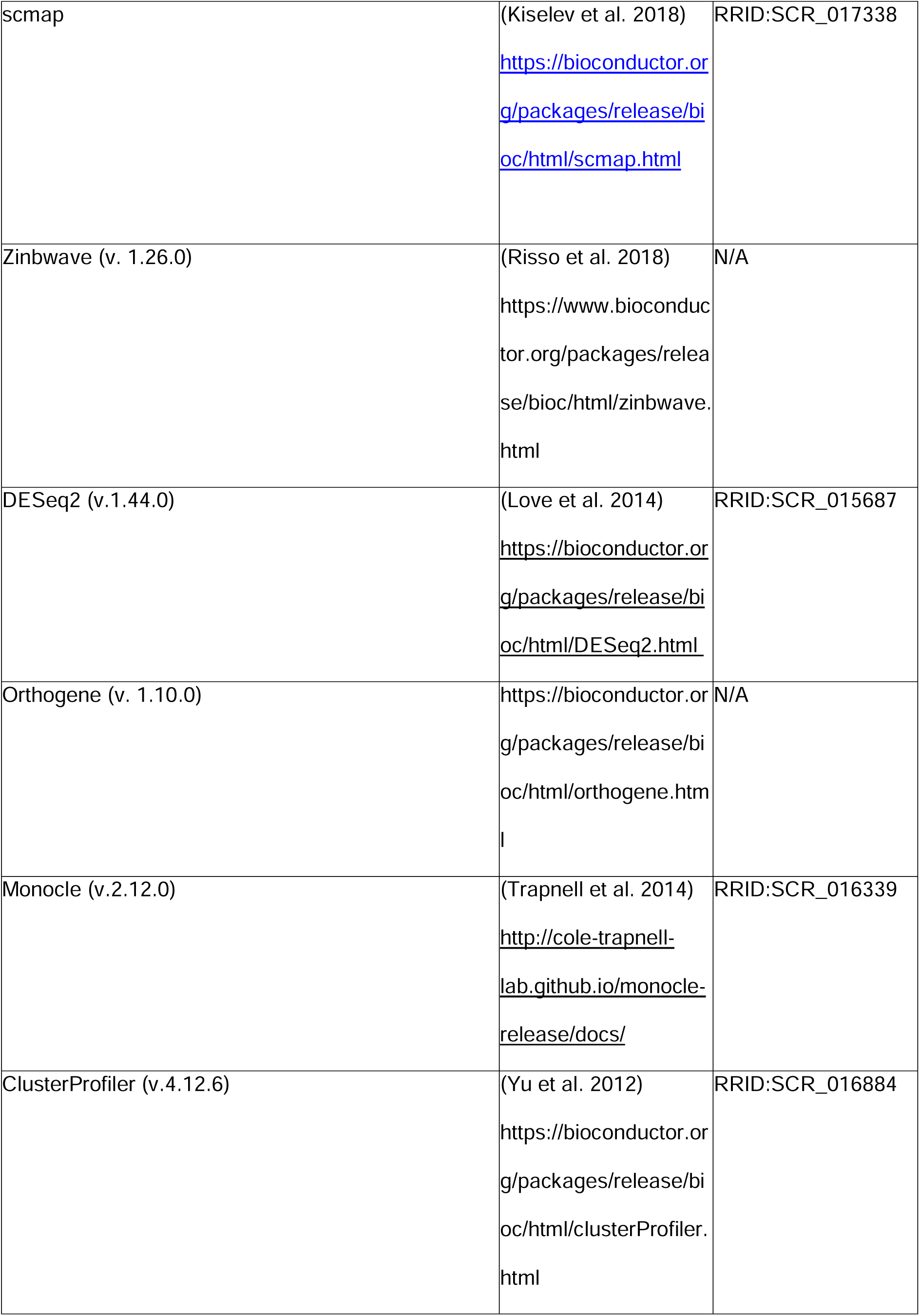
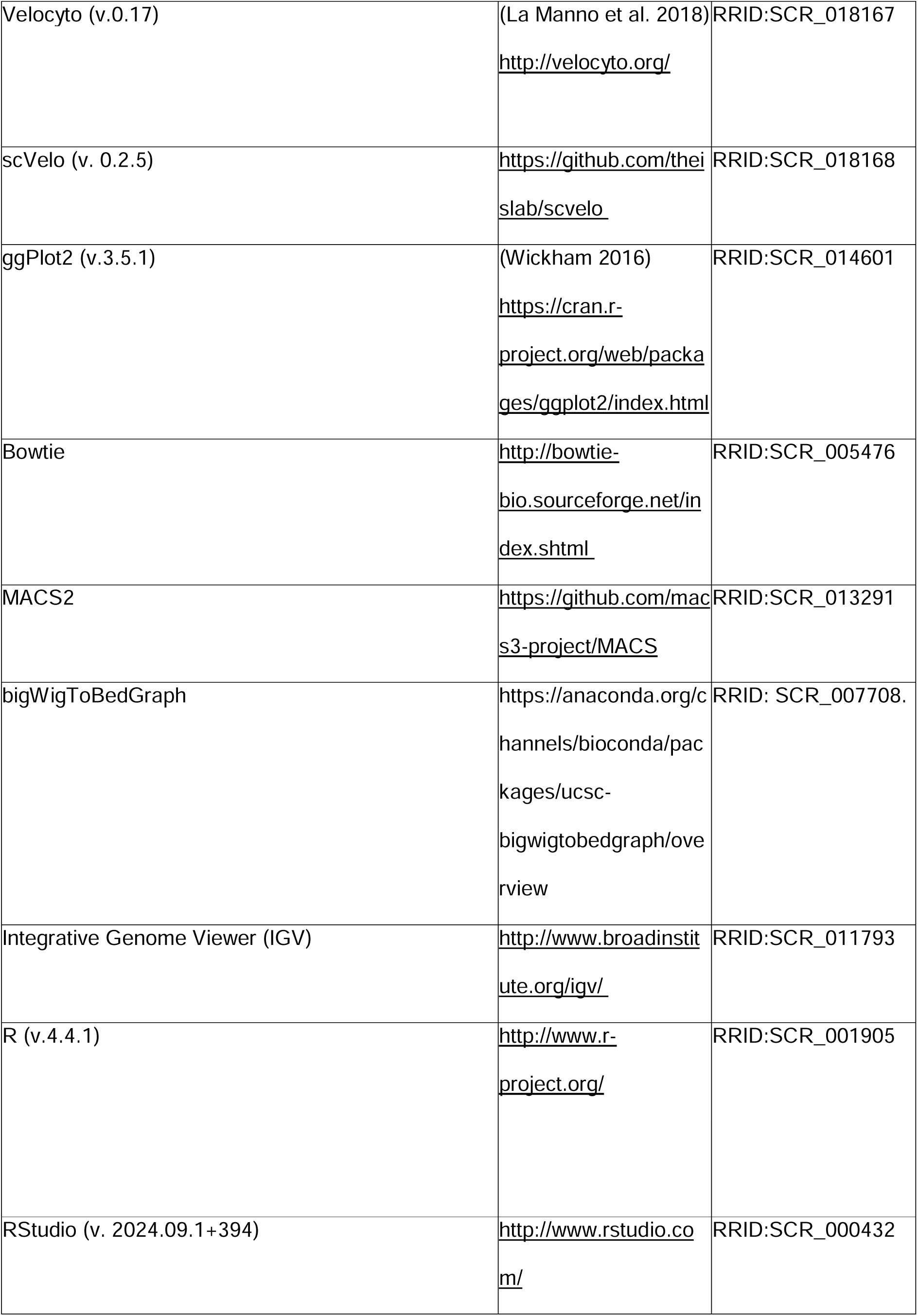
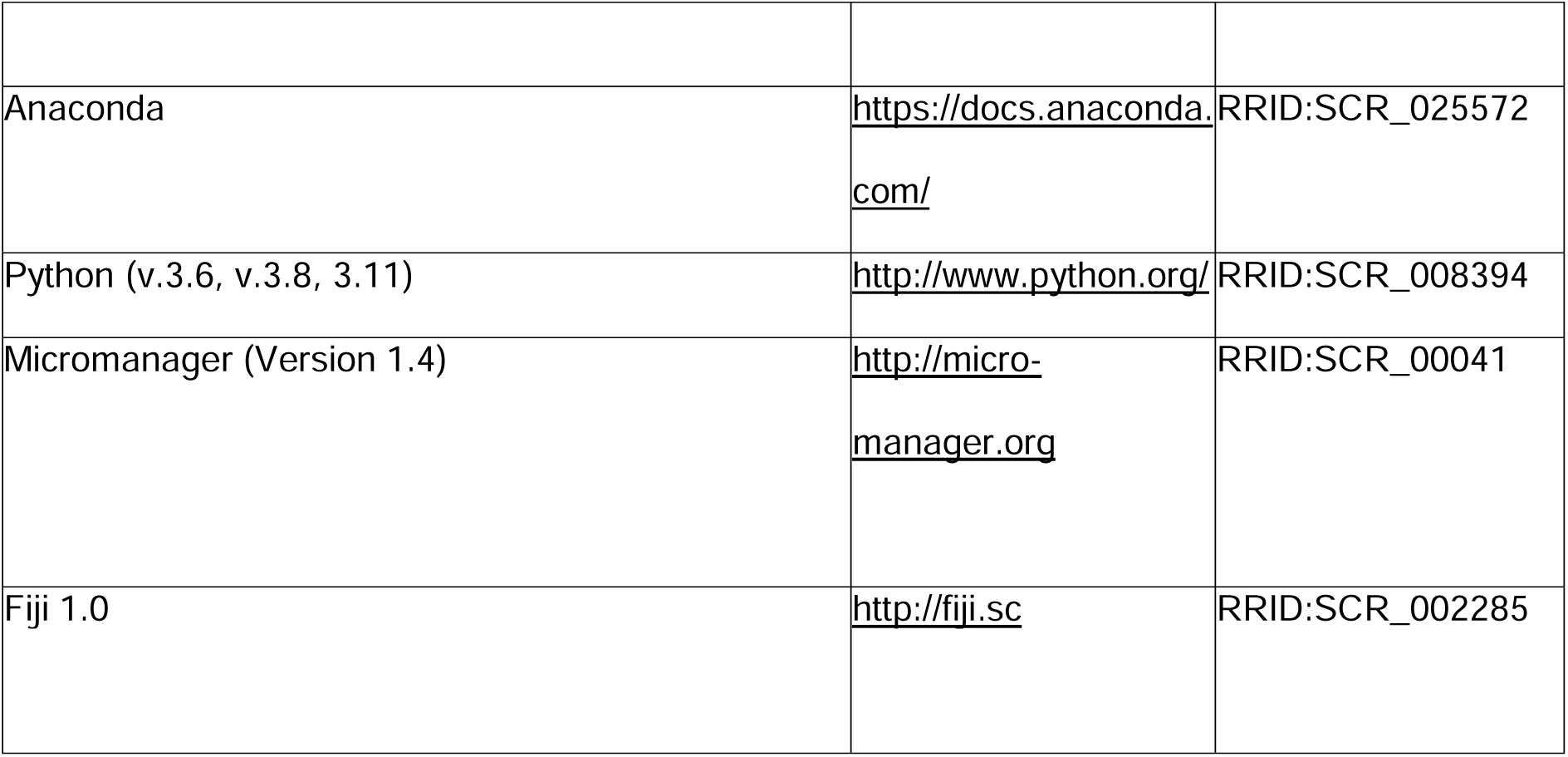
RESOURCES TABLE.

## RESOURCE AVAILABILITY

### Lead contact

Further information and requests for resources and reagents should be directed the Lead Contact, E. Elizabeth Patton.

### Materials availability

Newly generated materials from this study are available upon request to the Lead Contact, E. Elizabeth Patton.

### Data and code availability

scRNA-seq experiments have been submitted to GEO and will become available upon publication. All other data and codes supporting the findings of this study are available from the Lead contact upon reasonable request.

## EXPERIMENTAL MODELS

Zebrafish were maintained in accordance with UK Home Office regulations, UK Animals (Scientific Procedures) Act 1986, amended in 2013, and European Directive 2010/63/EU under project license 70/8000, P8F7F7E52 and PP7283023. All experiments were approved by the Home office and AWERB (University of Edinburgh Ethics Committee).

Fish stocks used were: wild-type AB, *mitfa^vc7^* and mitfa^w2^, here referred as *mitfa^nacre^* (Johnson et al. 2011; Zeng et al. 2015), *Tg(crestin:mCherry)* (Brombin et al. 2022; Brunsdon et al. 2022), *Tg(mitfa:GFP), Tg(Xla.Tubb:dsred) –* here referred as *Tg(NBT:DsRed)* (Dooley et al. 2013), *Tg(crestin:creERT2)* (generated for this study from the plasmid kindly provided from Charles Kaufman, Washington University), *Tg(ubi:loxP-GFP-loxP-mCherry; here referred as ubi:Switch*) (Mosimann et al. 2011). Combined transgenic and mutant lines were generated by crossing. Adult fish were maintained at ∼28.5°C under 14:10 light:dark cycles. Embryos and juveniles were kept at either 25°C, 28.5°C or 32°C and staged according to the reference table provided by Kimmel and colleagues (Kimmel et al. 1995) or Parichy and colleagues (Parichy et al. 2009).

## METHODS

### Generation of transgenic lines

The *p5E-crestin* promoter (Brombin et al., 2022), *pME-creERT2* (Travnickova et al., 2022) and the SV40 polyA sequence from *p3E-polyA* (Tol2Kit v1.2, plasmid #: 302) were assembled by multisite gateway assembly into a *pDestTol2CG2* destination vector (Tol2Kit v1.2, plasmid #: 395) to generate the *pEXPCG2(crestin:creERT2)* expression vector using the LR Clonase™ II Plus enzyme (Thermo Fisher) following standard protocols. The *pEXP* vector was mixed with *Tol2* mRNA (*in vitro* transcribed with the Ambion mMessage mMachine SP6 Kit, Thermo Fisher, from the Tol2Kit *pCS2FA-transposase* plasmid – plasmid #: 396); and microinjected into 1-cell stage either AB or ubi:Switch embryos, at a final concentration of 25 pg/nl and 35 pg/nl respectively. Zebrafish embryos expressing the *crestin:creERT2* transgene and GFP transgenic marker in the heart (myl7:eGFP is part of pEXPCG2 construct for transgene selection purposes) were selected and grown to adulthood before crossing with *ubi:Switch* and *ubi:Switch; mitfa^vc7/vc7^* zebrafish to obtain the F1 generation (Kwan et al. 2007; Roure et al. 2007).

### 4 – Hydroxytamoxifen (4-OHT) treatment and lineage tracing

*Tg(crestin:creERT2; ubi:Switch)* and *Tg(crestin:creERT2; ubi:Switch); mitfa^vc7/vc7^* fish were simultaneously pair mated. The resulting embryos were collected the following day and raised at 32°C until 24 hpf and then treated with 4-Hydroxytamoxifen (4-OHT, Sigma, H6278). 4-OHT stock was prepared by dissolving the powder in ethanol to reach 10 mM concentration, thoroughly vortexed and aliquoted into 100 µl aliquots. The 4-OHT stock solution was then diluted into 200 µM intermediate stock in E3, vortexed thoroughly while protected from light and then diluted in E3 to the final working concentration, 10 µM. Embryos were placed into six-well plates with 3 ml 10 µM 4-OHT or 0.1 % ethanol as a vehicle control, protected from the light and incubated at 32°C for 8h. Embryos were then washed three times with E3 and incubated in E3 at 32°C prior to imaging. Treated embryos of both genotypes were raised at 32°C for 5 weeks before being divided into 2 batches/genotype and raised for a further 5 weeks either at the permissive temperature for Mitfa expression (24°C) or at the non-permissive one (32°C). Only treated embryos displaying ‘switched’ mCherry^+^ signal in the McSCs at 4 dpf were raised beyond 5 dpf.

### Vibratome sections

4-OHT – treated *Tg(crestin:creERT2; ubi:Switch)* and *Tg(crestin:creERT2; ubi:Switch); mitfa^vc7/vc7^* juveniles were culled by overdose of anaesthetic (1 g l^−1^ MS-222, followed by death confirmation), cut in the middle of a clearly visible clone (previously determined using fluorescence stereoscope as described here below) before being fixed in 4% PFA overnight at 4°C. The fixed juvenile fish were then washed with PBS and embedded in 4% agarose. 200 µm sections were cut using a Leica VT1200S vibratome.

### Imaging

4-OHT – treated *Tg(crestin:creERT2; ubi:Switch)* and *Tg(crestin:creERT2; ubi:Switch); mitfa^vc7/vc7^* embryos were anaesthetised and screened under a Leica M205FCA stereoscope equipped with Leica K8 camera and operated via LASX software (Leica) using GFP and mCherry fluorescent filters. Embryos positive for the presence of mCherry signal in the McSCs at 4 dpf and were then raised as described previously. The same microscope was used to follow 4-OHT-treated embryos. Imaging was performed at around 5 weeks, at the end of metamorphosis or at terminal points (∼10 weeks). Larvae older than 4 dpf were first soaked in 5 mg/ml epinephrine (Sigma-Aldrich) for 5 minutes to contract the melanin within the differentiated melanocytes and anaesthetised prior to imaging. 10 week old fish were first soaked in 5 mg/ml epinephrine (Sigma-Aldrich) for 5 minutes and imaged in overdose of anaesthetic.

For confocal imaging, anaesthetised embryos were mounted laterally in low-melting point agarose in a glass-bottom 6-well-plate (Cellvis). Fish older than 5 dpf were first incubated in overdose of anaesthetic and 5mg/ml epinephrine (Sigma-Aldrich) for 10 minutes prior to mounting in low-melting point agarose and imaging. Vibratome sections were simply mounted in low-melting point agarose prior to imaging. Images of randomly picked embryos, selected adult fish or sections were acquired using a 4X/0.3, a 10X/0.5 or a 20X/0.75 lens on the multimodal Imaging Platform Dragonfly (Andor technologies, Belfast UK) equipped with 488 and 561 nm lasers built on a Nikon Eclipse Ti-E inverted microscope body with Perfect focus system (Nikon Instruments, Japan). Data were collected in Spinning Disk 40μm pinhole mode on the Zyla 4.2 sCMOS camera using a Bin of 1 and no frame averaging using Andor Fusion acquisition software.

Data were analysed using Fiji 1.0 and 64bit Java8. Representative of 3 biological repeats.

### Single cell suspensions, fluorescence activated cell sorting and library preparation

For consistency with the MITF^WT^ condition that included treatment with the ErbB-inhibitor (Brombin et al. 2022), we sequenced the transcriptome of cells from both untreated and ErbBi-treated MITF^OFF^ embryos, but excluded the latter from further analysis to concentrate on the expanded niche cells. 300 – 400 *Tg(crestin:mCherry; mitfa:GFP); mitfa^vc7/vc7^* embryos at 4 hpf were divided in two equally sized batches and arrayed in 6-well plates (Corning) containing either E3 or 5 μM AG1478 (ErbBi) in 3 ml of E3 till 24 hpf. A single cell suspension of each batch of embryos was then produced following the method described by Manoli and colleagues (Manoli and Driever 2012).

4-OHT – treated *Tg(crestin:creERT2; ubi:Switch)* and *Tg(crestin:creERT2; ubi:Switch); mitfa^vc7/vc7^* fish were first soaked in 5 mg/ml epinephrine (Sigma-Aldrich) for 5 minutes and imaged in overdose of anaesthetic as described above. Fish were then beheaded, and their intestine were dissected before cell dissociation as previously described (Travnickova et al. 2019).

Samples were sorted by a FACSAria2 SORP instrument (BD Biosciences UK). Green fluorescence was detected using GFP filters 525/50 BP and 488 nm laser, red fluorescence was detected using RFP filters 582/15 BP and 561 nm laser, and live cells selected with DAPI using DAPI filters 450/20 BP and 405 nm laser. Prior to sorting for fluorescence levels, single cells were isolated by sequentially gating cells according to their SSC-A vs. FSC-A and FSC-H vs FSC-W profiles following standard flow cytometry practices. Cells with high levels of DAPI staining were excluded as dead or damaged. Cells from wild-type stage matched embryos (without transgenes) were used as negative control to determine gates for detection of mCherry and/or GFP fluorescence. Then cells were purified according to these gates. We collected 20,000 GFP+, mCherry+ and or double positive cells from either treated or untreated *Tg(crestin:mCherry; mitfa:GFP); mitfa^vc7/vc7^ embryos.*For lineage tracing experiment, we collected 23,287 mCherry+ cells from *Tg(crestin:creERT2; ubi:Switch)* fish kept at 32°C for 10 weeks (MITF^WT^, Group 1); 10,274 mCherry+ cells from *Tg(crestin:creERT2; ubi:Switch)* fish kept at 32°C for 5 weeks and then downshifted to 24°C for another 5 weeks (MITF^WT^, Group 2); 28,000 mCherry+ cells from *Tg(crestin:creERT2; ubi:Switch); mitfa^vc7/vc7^* fish kept at 32°C for 10 weeks (MITF^OFF^, Group 3) and 31,191 mCherry+ cells from *Tg(crestin:creERT2; ubi:Switch); mitfa^vc7/vc7^* fish kept at 32°C for 5 weeks and then downshifted to 24°C for another 5 weeks (MITF^REGEN^, Group 4). Fluorescent cells were collected in 100µl of 0.04% Bovine Serum Albumin (BSA)/PBS in DNA-LoBind tubes (Fisher Scientific), spun down at 300G at 4°C, resuspended in 34 µl of 0.04% BSA/PBS, and immediately processed using the Chromium platform (10x Genomics) with one lane per sample. Single-cell mRNA libraries were prepared using either the single-cell 3’ solution V2 kit (10x Genomics) – embryos or the single-cell 3’ solution V3.1 kit (10x Genomics) – metamorphic fish. Quality control and quantification assays were performed using High Sensitivity DNA kits on a Bioanalyzer (Agilent).

Libraries from *Tg(crestin:mCherry; mitfa:GFP); mitfa^vc7/vc7^*embryos were sequenced on an Illumina NovaSeq platform (2 lanes of a S1 flowcell, read 1: 26 cycles, i7 Index: 8 cycles, read 2: 91 cycles). Libraries from 4-OHT – treated *Tg(crestin:creERT2; ubi:Switch)* and *Tg(crestin:creERT2; ubi:Switch); mitfa^vc7/vc7^* fish were sequenced on an Illumina NextSeq2000 platform (2 lanes of a P3 flowcell, 100 cycles). Each sample was sequenced to an average depth of at least 1750 million total reads. This resulted in an average read depth of ∼50,000 reads/cell after read-depth normalisation.

## QUANTIFICATION AND STATISTICAL ANALYSIS

### Bioinformatics analysis

#### scRNA-seq data processing and quality check

FastQ files were aligned using the CellRanger (v.2.1.1 or v.7.2.0 10x Genomics) pipeline to custom zebrafish STAR genome index using gene annotations from Ensembl GRCz11 release 94 with manually annotated entries for *GFP*, *mCherry*, *mitfa* intron 5 and *mitfa* intron 6 transcripts, filtered for protein-coding genes (with Cell Ranger *mkgtf* and *mkref* options). Final cellular barcodes and UMIs were determined using Cell Ranger. Libraries were aggregated (using 10X Cell Ranger pipeline ‘cellranger aggr’ option), with intermediary depth normalization to generate a gene-barcode matrix.

Gene-cell matrices (From *Tg(crestin:mCherry; mitfa:GFP); mitfa^vc7/vc7^*embryos total: 2748, untreated embryos: 1593, ErbBi-treated embryos: 1155; from 4-OHT – treated *Tg(crestin:creERT2; ubi:Switch)* and *Tg(crestin:creERT2; ubi:Switch); mitfa^vc7/vc7^* fish total: 3380; MITF^WT^, Group 1: 1016; MITF^WT^, Group 2: 342; MITF^OFF^, Group 3: 1044; MITF^REGEN^, Group 4: 978) were uploaded to R (v. 4.3.2) and standard quality control metrics performed using the Scater package (v.1.30.1) (McCarthy et al. 2017). Only cells with total features >700 (embryos) or >200 (adults), log10 total counts > 3.0, and mitochondrial gene counts (%) < 10 were considered as high quality and kept for further analyses (From *Tg(crestin:mCherry; mitfa:GFP); mitfa^vc7/vc7^*embryos total: 2081, untreated embryos: 1314, ErbBi-treated embryos: 767; from 4-OHT – treated *Tg(crestin:creERT2; ubi:switch)* and *Tg(crestin:creERT2; ubi:switch); mitfa^vc7/vc7^* fish total: 3257; MITF^WT^, Group 1: 1006; MITF^WT^, Group 2: 331; MITF^OFF^, Group 3: 953; MITF^REGEN^, Group 4: 967). Prediction of the cell cycle phase was performed using the cyclone function in the Scran (v.1.12.1) (Lun et al. 2016).

#### Clustering, UMAP visualisation and cluster calling

The Louvain clustering of the separated libraries was performed with Seurat (Hao et al. 2021) using the *FindNeighbors* and *FindClusters* functions (From *Tg(crestin:mCherry; mitfa:GFP); mitfa^vc7/vc7^* embryos, only untreated embryos selected for further analysis: dims= 12, resolution= 0.5; from 4-OHT – treated *Tg(crestin:creERt2; ubi:Switch)* and *Tg(crestin:creERt2; ubi:Switch); mitfa^vc7/vc7^* fish: dims= 11, resolution= 0.5) after performing linear dimensionality reduction and checking the dimensionalities of the datasets visualised with elbow plots. Data were projected onto 2 dimensional spaces using Uniform Manifold Approximation and Projection (UMAP) (McInnes et al. 2018) using the same dimensionality values listed above. Cluster specific genes were identified using the *FindAllMarkers* and *FindMarkers* function in Seurat (v.4.0.0) with default parameters (Wilcoxon Rank Sum test that compares a single cluster against the others). Cluster calling was done both manually by comparing to published marker genes for specific cell types and by making unbiased pairwise comparisons based on gene overdispersion against published datasets – GEO #: GSE112294 (Wagner et al. 2018), GEO #: GSE131136 (Saunders et al. 2019), NCBI SRA #: PRNJNA56410 (Farnsworth et al. 2020), GEO# GSE190115 (Frantz et al. 2023b) and GEO#: GSE178364 (Brombin et al. 2022) – using the scmap package (v.1.6.0) (Kiselev et al. 2018). Datasets were further subclustered after the first round of cluster calling to eliminate contaminants (**Supplementary Figure S1** and **S3**) using the same strategy described above. The new parameters were: From *Tg(crestin:mCherry; mitfa:GFP); mitfa^vc7/vc7^* untreated embryos: dims= 12, resolution= 0.4; from 4-OHT – treated *Tg(crestin:creERT2; ubi:Switch)* and *Tg(crestin:creERT2; ubi:Switch); mitfa^vc7/vc7^* fish: dims= 9, resolution= 0.5). Cycling status was determined as previously described (Sur et al. 2023). Integration of the embryonic originated datasets was performed following the Reciprocal Principal Component Analysis (RPCA) algorithm within the Seurat (v5.1.0) package (k.anchor = 12). Plots were generated either using Seurat (v.4.0.0, v. 5.0.1, v.5.1.0) (Hao et al. 2024) or ggplot2 (v.3.5.0) (Wickham 2016).

#### Differential expression and pathway analyses

The differential expression analyses (DEAs) were performed using the DeSeq2 package (v. 1.42.0 (Love et al. 2014)) after correction for zero-inflated matrix with Zinbwave using default parameters (v. 1.24.0 (Van den Berge et al. 2018)). Genes with log2 fold change >0 and p-value <0.05 were considered significantly upregulated. For direct comparison to human melanoma cell lines, human orthologous genes were determined with the orthogene package (v. 1.10.0, https://doi.org/doi:10.18129/B9.bioc.orthogene). Datasets used for comparison were GEO#: GSE178364 (Brombin et al. 2022) and GEO#: GSE163646 (Dilshat et al. 2021). Pathway analyses were performed using the ClusterProfiler R package (v.4.10.0). Pathway overrepresentation analyses were based on Gene Ontology-Biological Process database. Pathways with *P*.adj < 0.05 (hypergeometric test, BH adjusted, p-value cut off=0.05, q-value cut off=0.05) were considered to be significant and listed in **Supplementary Table S4**. The results were plotted using the ggplot2 package (v.3.5.0). Additional plots were generated with Seurat v. 5.1.0).

#### Pseudotime analyses and comparison of the developmental lineages

The standard pseudotime analysis (**Figure 3**) was performed as previously described (Brombin et al. 2022). The integrated pseudotime analysis comparing cells from *Tg(crestin:mCherry; mitfa:GFP); mitfa^vc7/vc7^* untreated embryos to the dataset from WT stage-matched embryos (GEO#: GSE178364 (Brombin et al. 2022)) (**Figure 4D-E**) was performed as follows. The top 1000 highly dispersed genes among the WT dataset were chosen as feature genes to reconstruct the developmental trajectory using the *setOrderingFilter*, *reduceDimension*, and *orderCells* functions of Monocle (v.2.12.0). We used the default parameters (except for max_components = 4 and norm_method = log) to generate the 3D trajectory during dimensionality reduction. The same set of genes was used to order the cells from the MITF^OFF^ embryos and then the combined results were plotted using the *PlotComplexTrajectories* function to highlight divergent states in the MITF^OFF^ dataset.

#### Analysis of the MITF ChIP-seq and CUT&RUN datasets

Publicly available MITF ChIP–seq and CUT&RUN datasets were retrieved from the GEO repository (accessions GSE61965, GSE153020, and GSE267956) (Laurette et al. 2015; Kenny et al. 2022; Vu et al. 2024). Raw sequencing reads were aligned to the human reference genome (hg38) using Bowtie. Peaks were identified with MACS2 (p < 1×10⁻L) using matched input DNA or IgG controls. Signal intensities were generated as WIG tracks and converted to BEDGRAPH format with bigWigToBedGraph. Processed data were visualized and peak profiles inspected using IGV.

#### RNA velocity analyses

RNA velocity analyses were performed with the Velocyto (v.0.17.16) (La Manno et al. 2018) and scVelo (v. 0.2.5) (Bergen et al. 2020) Python packages. Velocyto was used to generate loom files for each experimental condition, according to package instructions. Velocity estimations were computed using scVelo’s dynamical modelling approach and visualised as streamlines over UMAP.

## SUPPLEMENTAL INFORMATION

### Supplementary Figure Legends

**Figure S1.** Analysis pipeline, quality check results and cluster calling of the embryonic dataset.

**Figure S2.** Flow cytometry plots with delineated gating strategy for lineage tracing scRNA-seq

**Figure S3.** Quality check results and cluster calling for the lineage tracing-related dataset.

**Figure S4.** Proportions of expressed genes and cells in clusters and cycling scores in the lineage tracing-related dataset.

### Supplementary Tables Legends

**Table S1:** Metrics, clustering information, and cell states for all cells (Untreated *mitfa^vc7/vc7^* embryos).

**Table S2:** Metrics, clustering information, and cell states for all cells (ErbBi-treated *mitfa^vc7/vc7^* embryos).

**Table S3:** Cluster markers (Untreated *mitfa^vc7/vc7^* embryos). Comparison between indicated cluster relative to pooled cells of all other clusters.

**Table S4:** Common overrepresented GO terms between Neural Crest – Neuronal lineage and MITF^KO^ SK-MEL-28.

**Table S5:** Differential expression analysis of MIX+ (WT) vs *sox4a*^+^ neural crest (*mitfa^vc7^*

**Table S6:** Metrics, clustering information, and cell states for all cells (Lineage Tracing).

**Table S7:** Cluster markers (Lineage Tracing). Comparison between indicated cluster relative to pooled cells of all other clusters.

**Figure S1.** Analysis pipeline, quality check results and cluster calling of the embryonic dataset. **A.** Summary of the analysis pipeline followed **B – D.** Bar plots showing the relation to the Log_10_ library size **(B)**, the number of expressed genes (**C**), and the percentage of mitochondrial genes (**D**) of the aggregated libraries from cells pre-quality check (QC). **E – F.** Scatter plots distribution of the total features (genes) expressed in relation to the Log_10_ total counts (**E**) and the percentage of mitochondrial genes in relation Log_10_ total counts (**F**) of the aggregated libraries from embryos pre-QC. Red dashed lines indicate the QC thresholds (Total features by counts > 200; Log_10_ total counts > 3.0; Percentage of mitochondrial genes <10%). Gray areas highlight the cells that were considered low quality and excluded from downstream analyses. Orange: cells from untreated embryos, Blue: cells from ErbBi-treated embryos. **G.** Principal component analysis (PCA) plot of the aggregated libraries from cells post-QC. Orange: cells from untreated embryos, Blue: cells from ErbBi-treated embryos. **H.** Pie chart of the proportion of cells in a predicted cell cycle phase within the aggregated libraries from cells post-QC. **I.** UMAP of cells (n = 1314 cells) obtained from untreated 24 hpf zebrafish MITF^OFF^ embryos after Louvain clustering (dims= 12, resolution = 0.5) showing 10 clusters. Cells of the pigment lineage made up the bulk of the dataset (dashed circle). Contaminants are highlighted and were excluded in subsequent analyses. **J.** UMAP cells (n = 1111 cells) obtained from 24 hpf zebrafish MITF^OFF^ embryos after Louvain clustering (dims= 12, resolution = 0.4) after discarding the contaminant cell types shown in **I**. The novel cell populations are identified here with a dashed line. **K.** UMAP representations based on the scmap results obtained by comparison of the MITF^OFF^ dataset and the Wagner *et al*., 24 hpf dataset, the Farnsworth *et al*., 24 hpf dataset, and the Saunders *et al*., *sox10:cre+* 5 dpf zebrafish embryos dataset.. The “sox4a+ Neural crest” population (dotted line) failed to map consistently to any of the known populations. See also Figure 2G.

**Figure S2.** Flow cytometry plots with delineated gating strategy for lineage tracing scRNA-seq. A. FACS plot of wild type AB fish. B. FACS plot of *mitfa^vc7/vc7^* fish. C. FACS plot of an untreated *Tg(crestin:creERT2;ubi:Switch)* fish (GFP ‘unswitched’ cells). D. FACS plot of an untreated *Tg(crestin:creERT2;ubi:Switch)*; *mitfa^vc7/vc7^* (GFP ‘unswitched’ cells). E. FACS plot of a *Tg(crestin:creERT2;ubi:Switch)* fish treated with 10 µM 4-OHT for 8 hours and kept at 32°C for 10 weeks resulting in mCherry+ ‘switched’ cells. F. FACS plot of a *Tg(crestin:creERT2;ubi:Switch)*; *mitfa^vc7/vc7^* fish treated with 10 µM 4-OHT for 8 hours and kept at 32°C (MITF^OFF^) for 10 weeks resulting in mCherry+ ‘switched’ cells. G. FACS plot of a *Tg(crestin:creERT2;ubi:Switch)* fish treated with 10 µM 4-OHT for 8 hours and kept at 32°C for 5 weeks before being downshifted to 24°C for a further 5 weeks, resulting in mCherry+ ‘switched’ cells. H. FACS plot of a *Tg(crestin:creERT2;ubi:Switch)*; *mitfa^vc7/vc7^* fish treated with 10 µM 4-OHT for 8 hours and kept at 32°C for 5 weeks before being downshifted to 24°C for a further 5 weeks (MITF^REGEN^), resulting in mCherry+ ‘switched’ cells.

**Figure S3.** Quality check results and cluster calling for the lineage tracing-related dataset. **A.** Scatter plots of the total features (genes) expressed in relation to the Log_10_ total counts (**Left panel**) and the percentage of mitochondrial genes in relation Log_10_ total counts (**Mid panel**) of the aggregated libraries pre-QC. Red dashed lines indicate the QC thresholds (Total features by counts > 200; Log_10_ total counts > 3.0; Percentage of mitochondrial genes <30%). Gray areas highlight the cells that were considered low quality and excluded from downstream analyses. Green: cells from *Tg(crestin:creERt2; ubi:Switch)* fish kept at 32°C (Group 1), Red: cells from *Tg(crestin:cre^ERt2^; ubi:Switch)* fish raised at 32°C and downshifted to 24°C (Group 2); Orange: cells from *Tg(crestin:creERt2; ubi:Switch); mitfa^vc7/vc7^* fish kept at 32°C (Group 3), Blue: cells from *Tg(crestin:creERt2; ubi:Switch); mitfa^vc7/vc7^* fish kept at 32°C (Group 4). **B.** Principal component analysis (PCA) plot of the aggregated libraries from cells post-QC. Legend is shared with **A**. **C.** UMAP of mCherry+/DAPI− cells (n = 3257 cells) obtained from 16 fish after Louvain clustering (dims= 11, resolution = 0.5) showing 17 clusters. Cells of the pigment lineage made up the bulk of the dataset. Contaminants are highlighted and were excluded in subsequent analyses. **D.** UMAP of mCherry^+^ /DAPI− cells (n = 2660 cells) obtained from 4-OHT-treated fish after Louvain clustering (dims= 12, resolution = 0.4) and after discarding the contaminant cells shown in **C**. **E.** UMAP representations based on the scmap results generated by comparison of the dataset obtained from 4-OHT treated fish and the Frantz *et al*., regenerating fish dataset and the Saunders *et al*., *sox10:cre+* 1 mpf euthyroid fish dataset.

**Figure S4.** Proportions of expressed genes in the lineage tracing-related dataset. **A.** UMAP representation of Figure 6A, with colour change from gray (negative) to red (positive) based on the cycling status calculated as described in (Sur et al., 2023). **B.** Distribution of the expressed genes (Counts, **left**) and the Unique Molecular Identifiers (UMIs, **right**) in each cell of the dataset in the different clusters. **C.** UMAP in Figure 6A, split by experimental groups (MITF^WT^, MITF^OFF^, MITF ^REGEN^) for visualisation of the cell cluster distributions.

