## Supplementary Figures for "Loss of MITF activity leads to emergent cell states from the melanocyte stem cell lineage"

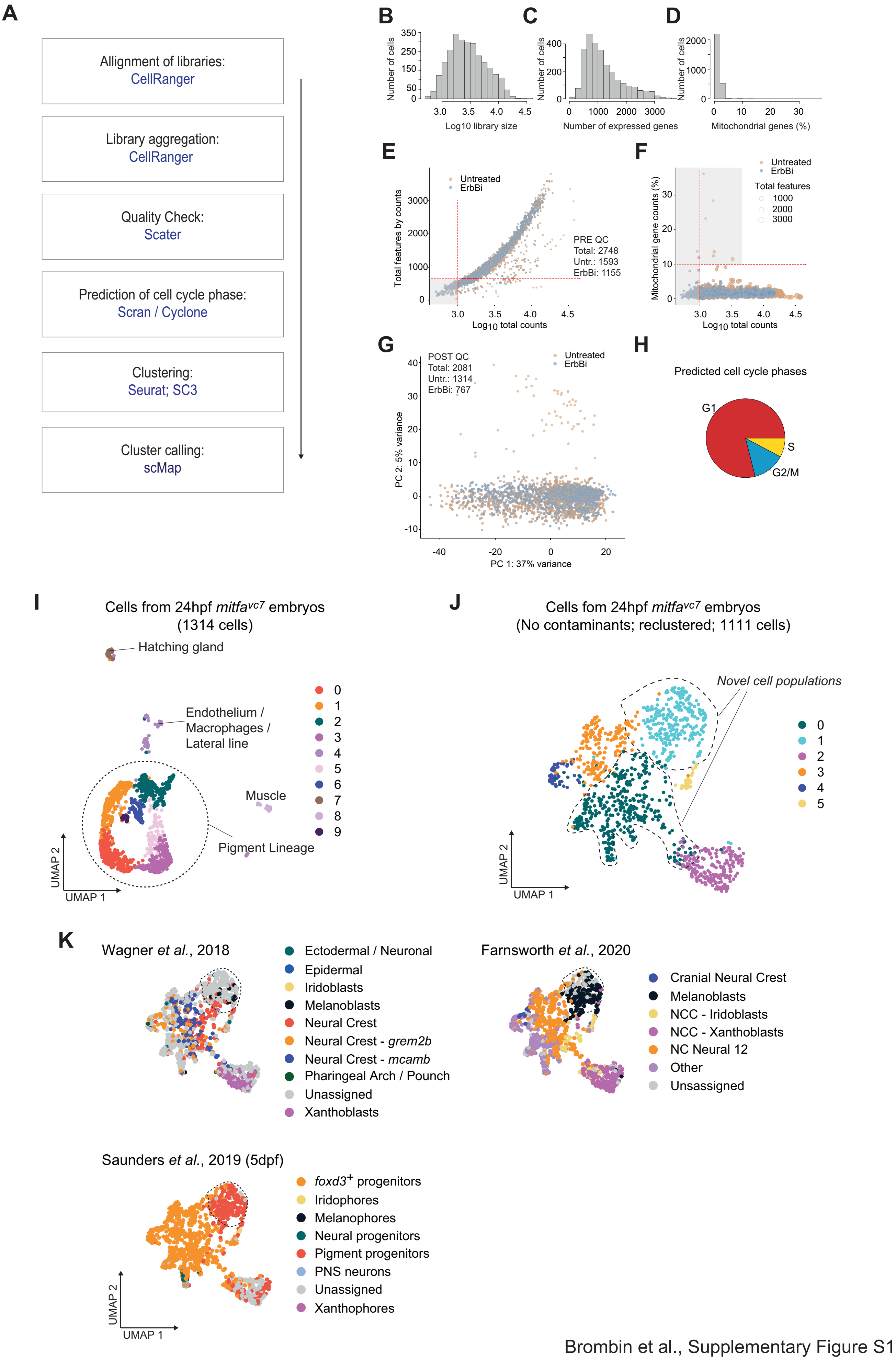


Brombin et al., Supplementary Figure S1

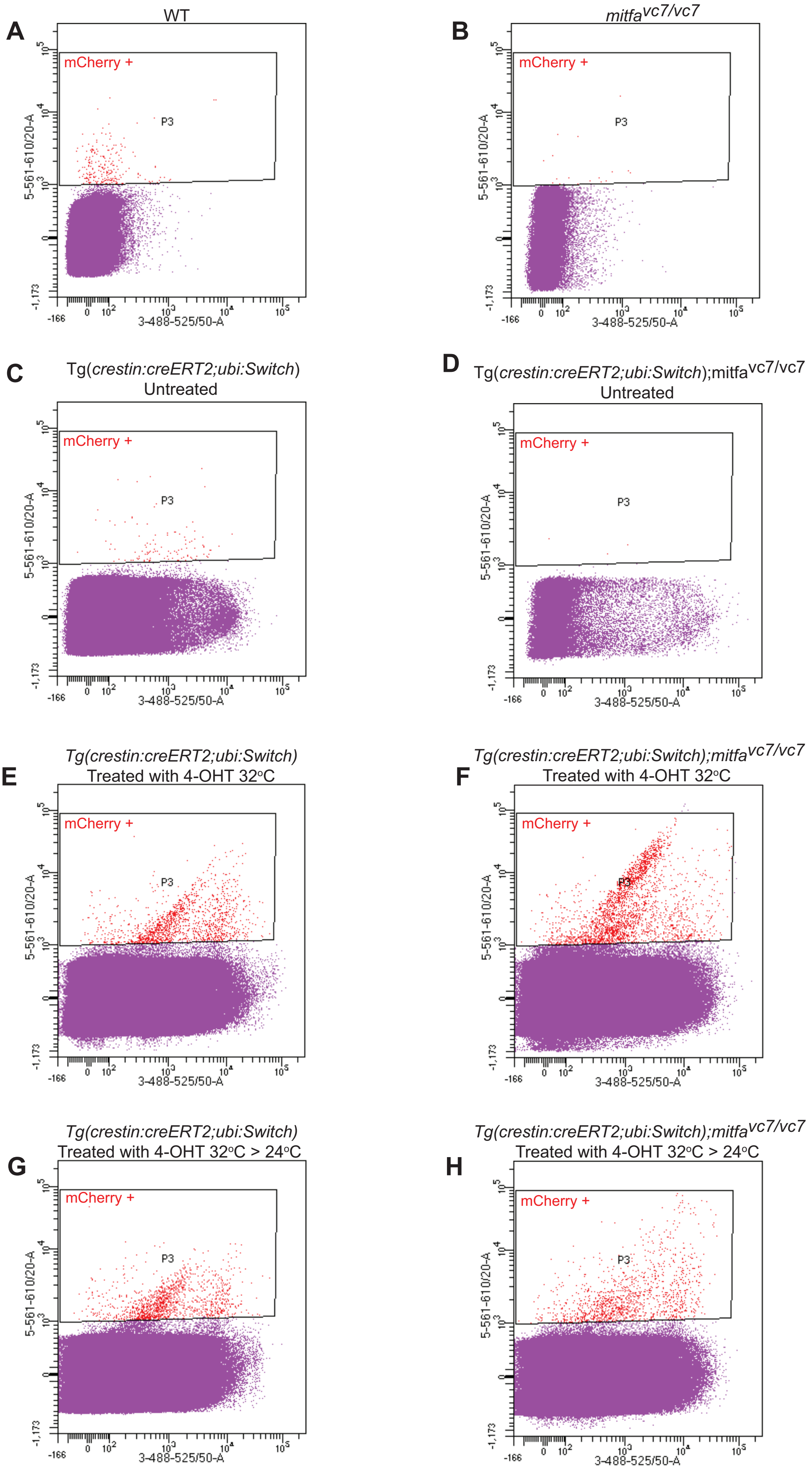


Brombin et al., Supplementary Figure S2

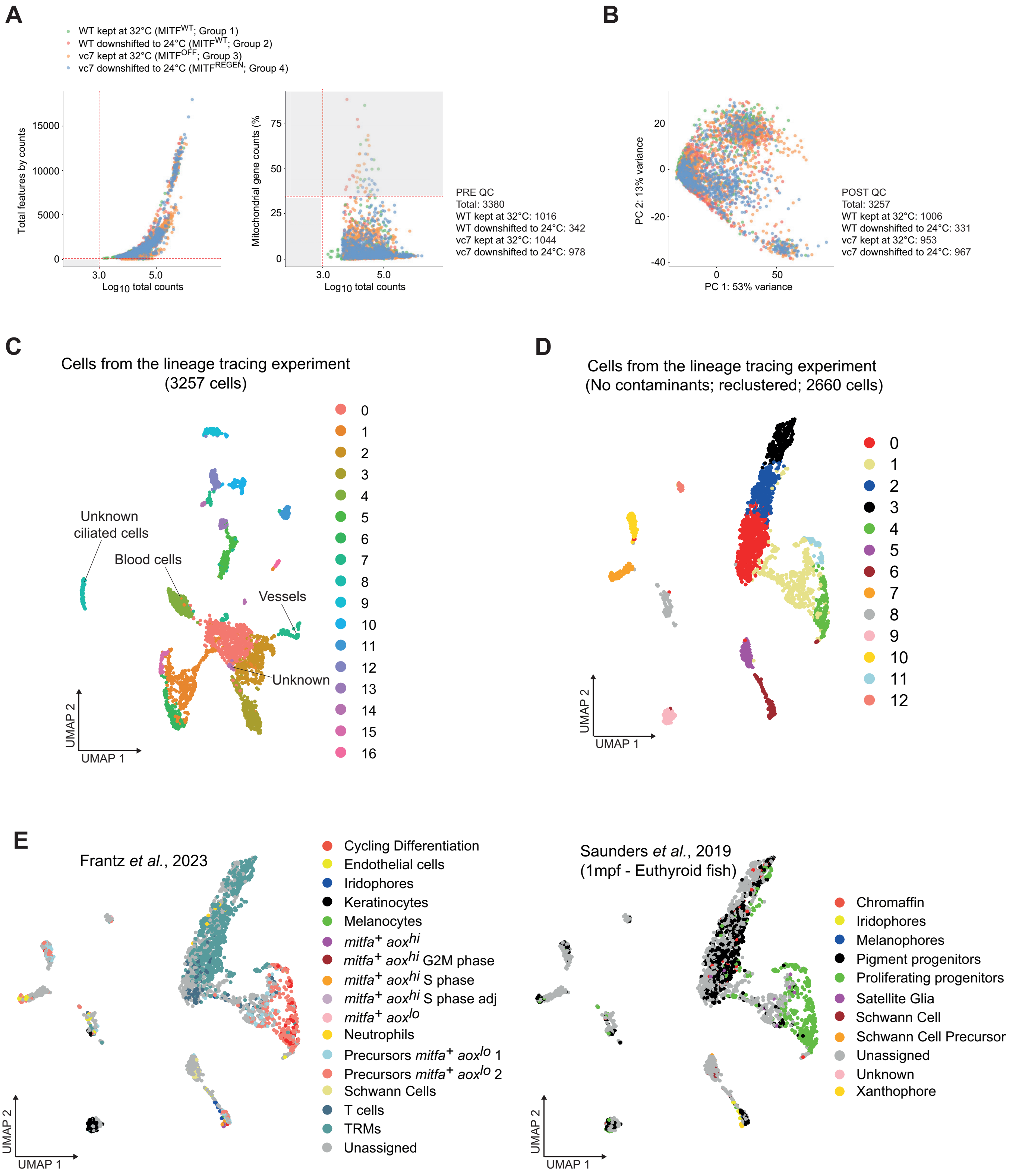


Brombin et al., Supplementary Figure S3

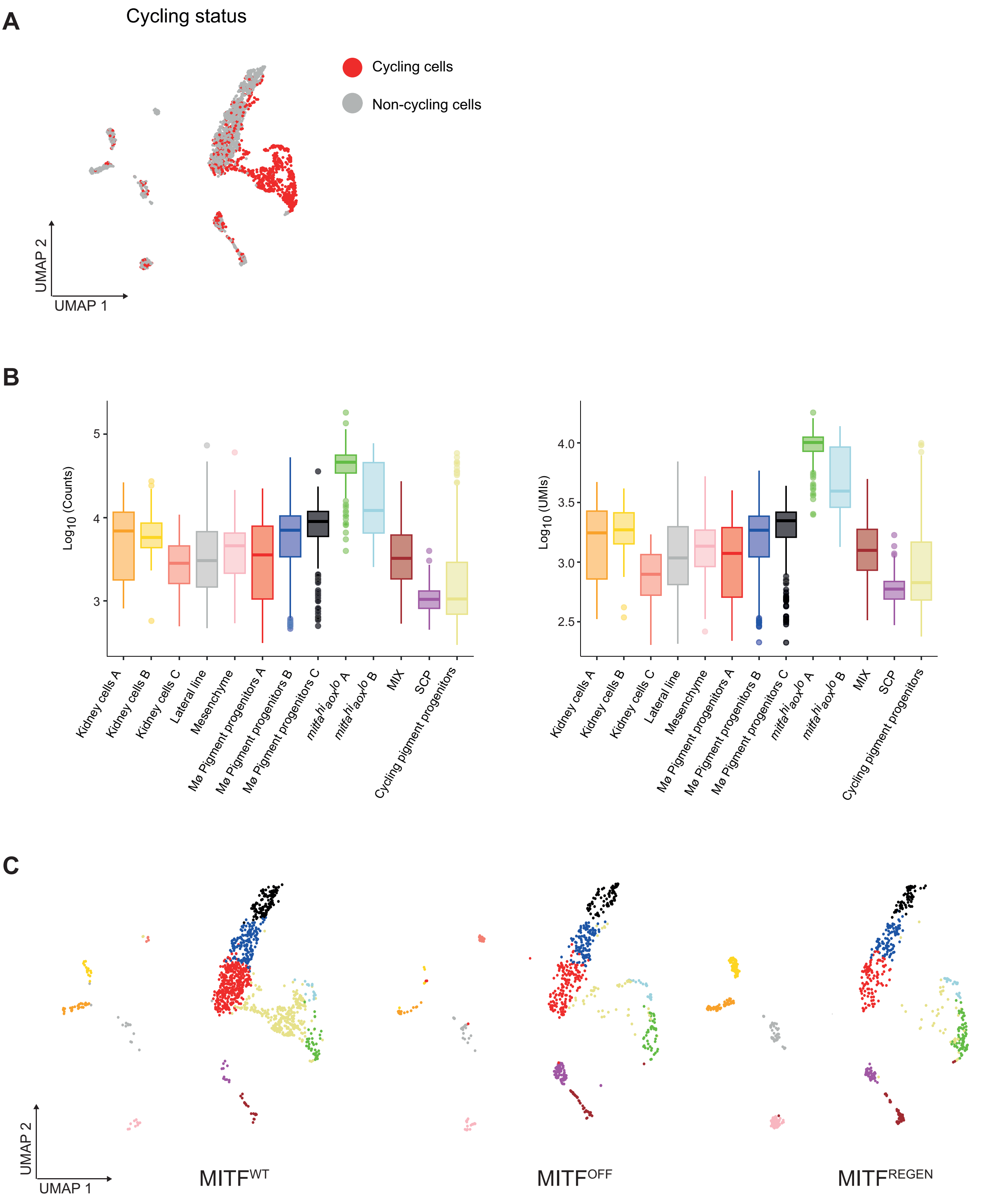


Brombin et al., - Supplementary Figure S4
